# Selective Pyroptosis in NF1-Deficient Cells through PKCδ Agonism

**DOI:** 10.64898/2026.05.19.726276

**Authors:** Liang Hu, Yuting Tang, Yuan Lin, Jay Pundavela, Katherine Chaney Scheffer, Abby Schaeper, Tilat Rizvi, Jianqiang Wu, Yi Zheng, Nancy Ratner

## Abstract

Pyroptosis, a lytic and immunogenic form of cell death, holds broad therapeutic potential, yet its selective induction in specific cell populations remains a fundamental challenge. Loss of the *NF1* tumor suppressor, one of the most frequent events across pediatric and adult cancers, elevates RAS-GTP and drives tumorigenesis through hyperactivated RAS signaling. Here we demonstrate that protein kinase Cδ (PKCδ) agonism selectively triggers pyroptosis in NF1-deficient cells by exploiting their dependency on KRAS. PKCδ directly phosphorylates KRAS at S39 and S181, inducing KRAS-GDP accumulation and driving endoplasmic reticulum translocation. The dually phosphorylated KRAS-GDP interacts with caspase-8 and competitively displaces inhibitory BCL2, promoting caspase-8/caspase-3/gasdermin-E-mediated pyroptosis. This vulnerability is conserved across multiple NF1-deficient tumor types, and PKC agonism suppresses NF1-deficient neurofibroma and malignant peripheral nerve sheath tumor growth *in vivo*. These findings establish the inactive KRAS-GDP as a functionally active signaling molecule and PKCδ agonism as a selective therapeutic strategy for NF1-deficient cancers.

## Main

Pyroptosis is a lytic form of regulated cell death mediated by gasdermin pore-forming proteins.^1^ Dysregulation of pyroptosis is implicated in diverse pathological conditions, including malignancies, infectious diseases, inflammatory disorders, and metabolic diseases.^1^ Unlike the well-known immunologically silent apoptosis, pyroptosis triggers inflammatory responses that activate anti-tumor immunity, positioning it as a powerful therapeutic modality.^2–4^ However, realizing this therapeutic potential faces a fundamental challenge because current pyroptosis - inducing agents lack cellular selectivity and cause indiscriminate tissue damage.^5^ This toxicity reflects the widespread expression of gasdermin proteins (e.g., GSDMD and GSDME) in healthy cells, where these executioner proteins are activated unselectively by therapeutic agents.^1, 5^ Achieving selective pyroptosis, particularly in cancer cells while sparing normal tissue, thus represents a critical unmet need across multiple disease contexts. Although disease-specific genetic alterations could enable selective targeting, pharmacological strategies to exploit such vulnerabilities for inducing selective pyroptosis remain largely unexplored.

*NF1* is a major tumor suppressor gene, ranking among the top five somatically mutated genes in pediatric cancers and among the 15 most frequently mutated genes in adult cancers.^6, 7^ Germline *NF1* mutations cause neurofibromatosis type 1, a tumor predisposition syndrome characterized by peripheral nerve sheath tumors, including benign neurofibromas and malignant peripheral nerve sheath tumors (MPNSTs).^8^ Individuals with neurofibromatosis type 1 also face increased risk of various other tumors, including breast cancer, lung cancer, and leukemia.^8, 9^ Current therapeutic options remain limited: MEK inhibitors for neurofibromas require long-term administration due to their cytostatic rather than cytotoxic effects,^10, 11^ and surgical resection and chemoradiotherapy for MPNSTs rarely achieve durable responses.^11–13^ Importantly, sporadic *NF1* mutation in other tumor types confers resistance to conventional therapies and significantly reduces patient survival.^14–17^ This paucity of effective treatments for diverse NF1-deficient cancers underscores an urgent need for novel therapeutic strategies.

*NF1* encodes neurofibromin, a RAS guanosine triphosphatase (GTPase)-activating protein (RasGAP) that accelerates the conversion of active GTP-bound RAS (RAS-GTP) into inactive GDP-bound RAS (RAS-GDP).^8, 18^ The current RAS paradigm holds that RAS-GDP is inactive owing to a lack of interaction with effectors, while RAS-GTP actively engages downstream effector.^18^ NF1 regulates six homologous RAS paralogs, including classical (H-/N-/K-RAS) and non-classical (R-/M-RAS, TC21) proteins.^19^ Consequently, NF1 loss increases RAS-GTP levels and enhances RAS signaling through the MEK/ERK and PI3K/AKT pathways, which are believed to drive NF1-associated tumorigenesis.^8^ Although TC21 has been implicated in the migration of NF1-deficient cells,^20^ the specific RAS paralog(s) driving their tumorigenesis and growth remain undefined—a gap that limits the identification of precise therapeutic targets for NF1-deficient tumors.

Protein kinase C (PKC) comprises a family of serine/threonine kinases, including nine isozymes categorized into three subfamilies: conventional (cPKC: α, β, γ), novel (nPKC: δ, ε, η, θ), and atypical (aPKC: ζ, ι). ^21, 22^ PKCs regulate multiple biological processes, including cell death and proliferation, and PKC dysfunction is implicated in many diseases, including cancer.^22^ While PKCs were initially considered oncogenes, the discovery that most cancer-associated PKC mutations cause loss-of-function suggests that many, if not all, PKC isozymes function as tumor suppressors.^21^ Supporting this tumor-suppressive role, PKC agonists have demonstrated therapeutic efficacy against human actinic keratosis,^23^ a precancerous skin condition, and in preclinical models of human colon and pancreatic cancers.^24, 25^ However, because tested PKC agonists activate multiple PKC isozymes, it remains unclear which specific isozyme(s) mediate antitumor effects and what molecular contexts determine their efficacy.

Here, we utilized NF1-deficient Schwann cells (SCs), the tumorigenic cells in neurofibromatosis type 1,^8^ as a model system and identified KRAS as essential for the tumorigenesis and growth of NF1-deficient SCs. Using prostratin as a tool compound, we demonstrate that PKC agonism selectively induces pyroptosis in NF1-deficient but not wild-type SCs, a vulnerability conferred by KRAS dependency and shared across multiple NF1-deficient cancer types. Mechanistically, PKCδ is the key PKC isozyme driving CASP8/GSDME-dependent pyroptosis through direct phosphorylation of KRAS at S39 and S181. KRAS-pS39 primarily upregulates KRAS-GDP levels by promoting KRAS-GTP hydrolysis and impeding KRAS-GDP dissociation, while KRAS-pS181 primarily drives endoplasmic reticulum translocation. The resulting dual-phosphorylated KRAS-GDP interacts with CASP8, competitively displacing BCL2 to promote CASP8 activation. Furthermore, PKC agonism via prostratin effectively inhibits NF1-deficient tumor growth in multiple preclinical models. These findings demonstrate that selective pyroptosis can be induced by a small molecule in a cell type-specific manner, define KRAS-GDP as a functional signaling molecule, and establish PKCδ agonism as a therapeutic strategy for NF1-deficient tumors.

## Results

### KRAS-dependency of NF1-deficient Schwann cells

Schwann cells (SCs) are glial cells of peripheral nervous system. NF1 loss in the SC lineage drives neurofibroma and MPNST formation.^26^ To model NF1 loss in a tumorigenic setting, we generated *NF1* knock-out (*NF1* KO) isogenic cell lines from immortalized human SCs (iHSC1λ)^27^ using CRISPR (**Figures S1A-S1C**). *NF1* KO iHSC1λ cells exhibited increased proliferation compared to wild-type (WT) control cells (**Figures S1D-S1E**) and displayed enhanced and sustained activation of RAS effectors (phosphorylated ERK and AKT; pERK, pAKT) upon serum stimulation (**Figure S1F**).

NF1 is a large protein that contains multiple functional domains, including a RasGAP domain that inactivates RAS.^8^ To test whether the loss of RasGAP activity is sufficient to drive these effects observed in NF1-deficient iHSC1λ cells, we transgenically expressed either green fluorescent protein (GFP), WT NF1 (NF1-WT), or a RasGAP-defective NF1 mutant (NF1-R1276P)^28^ in *NF1* KO iHSC1λ (**Figure S2A**). The NF1-R1276P cells, akin to the GFP cells, showed aberrant activation of RAS downstream effectors (pERK, pAKT) compared to the NF1-WT cells (**Figure S2B**), confirming that the mutant is inactive. Additionally, the NF1-R1276P and GFP cells showed similar growth kinetics, both growing faster than the NF1-WT expressing cells (**Figure S2C**). Soft-agar assays, a functional readout for *in vitro* tumorigenesis, revealed that the NF1-R1276P and GFP cells, but not the NF1-WT cells, formed colonies (**Figure S2D**). Collectively, these data indicate that the loss of RasGAP activity in NF1 drives the NF1-loss-related hyperproliferation and tumorigenesis *in vitro*.

To define the specific RAS paralog(s) mediating NF1-loss-related effects, we analyzed RAS paralog activation profiles in iHSC1λ cells. RAS pulldown assays using RAS paralog-specific antibodies in WT and *NF1* KO iHSC1λ cells showed that, among the six NF1-related RAS paralogs, only HRAS and KRAS were more activated after stimulation (**Figure 1A**). These two RAS paralogs also showed enhanced activity (higher GTP loading) in *NF1* KO iHSC1λ compared to WT iHSC1λ (**Figure 1A**). To characterize the roles of HRAS and KRAS in WT and *NF1* KO iHSC1λ cell proliferation, we first employed a dominant-negative approach. Three well-established dominant-negative RAS mutants (all with the S17N mutation) that differentially inhibit H-/N-/K-RAS paralogs^29^ (**Figure S3A**) were used. In *NF1* KO iHSC1λ cells, both dnHRAS and dnKRAS, but not dnNRAS, suppressed proliferation (**Figures S3B-S3C**). Despite most HRAS activity remaining unaffected by dnKRAS (**Figure S3A**), the comparable inhibitory effect of dnHRAS and dnKRAS on *NF1* KO iHSC1λ implied that HRAS is unlikely to be the key RAS paralog; instead, KRAS, the other active RAS paralog in iHSC1λ cells (**Figure 1A**), is likely essential for *NF1* KO iHSC1λ cell proliferation. Similar proliferation assays in WT iHSC1λ cells showed that both dnHRAS and dnNRAS effectively inhibited proliferation, whereas dnKRAS exhibited much weaker inhibitory effects (**Figures S3D-S3E**). Given that dnNRAS inhibits both NRAS and HRAS (**Figure S3A**), but NRAS was not activated in iHSC1λ cells (**Figure 1A**), the data suggest that HRAS is the critical RAS paralog mediating WT iHSC1λ cell proliferation. Collectively, these data imply that the proliferation of NF1-deficient SCs predominantly depends on KRAS, while wild-type SCs largely rely on HRAS.

**Figure 1.**
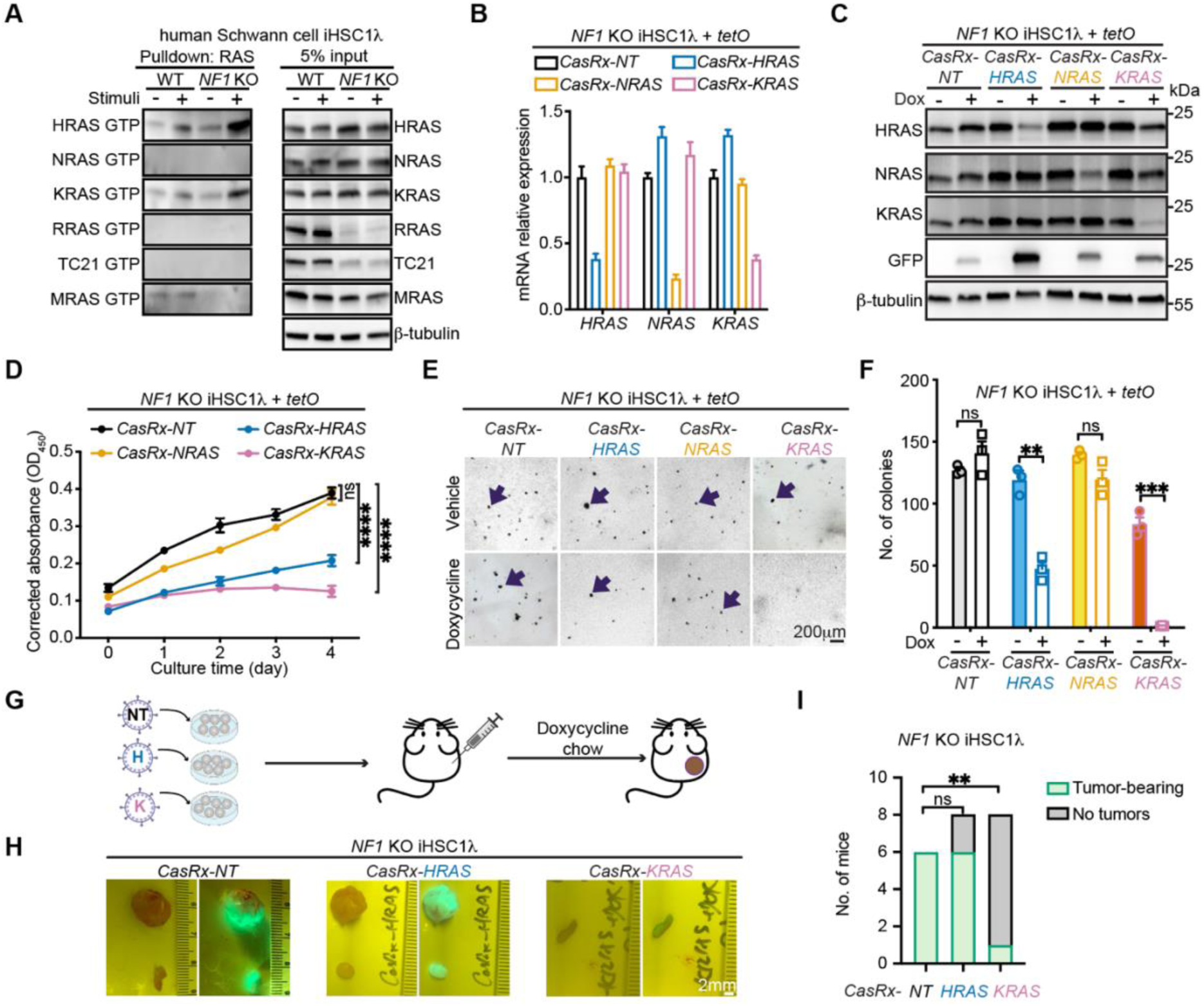
KRAS is critical for proliferation and tumorigenesis of NF1-deficient SCs. (**A**) Detection of active RAS-GTP (pull-down) and total RAS (input) by RAS pull-down assays (n=4). Stimuli: 15% serum plus β-heregulin, a Schwann cell growth factor. (**B-C**) *H-/N-/K-RAS* paralog-specific knockdown in *NF1* KO iHSC1λ cells using CasRx: qPCR analysis of mRNA levels (B) and immunoblotting analysis (C). NT: non-target; Dox: doxycycline (1μg/ml). (**D-F**) Analysis of *NF1* KO iHSC1λ cells upon knockdown of individual RAS paralog or NT: Growth kinetics (D) and soft-agar assays (E-F). Representative images (E) and colony quantification per well (F) of soft-agar assays. Arrows indicate formed colonies. (**G-I**) Xenograft analysis of *NF1* KO iHSC1λ cells with RAS paralog knockdown: experimental design (G), representative images of tumors under bright and fluorescence fields (H), and quantification of tumor formation(I). *CasRx*-NT: n=6; *CasRx-H/K-RAS*: n=8. Tumor threshold: dimension >5 mm. ns(non-significant), p>0.05; ∗p<0.05, ∗∗p<0.01, ∗∗∗p<0.001, ∗∗∗∗p<0.0001 by two-way ANOVA (D), t-test(F), or Fisher’s exact test (I). Scale bars are as indicated.

To confirm these findings, we established an inducible CasRx system^30^ to knock down *H/N/K-RAS* individually in both WT and *NF1* KO iHSC1λ cells (**Figures 1B-1C and S3F**). In *NF1* KO iHSC1λ cells, knockdown of either *HRAS* or *KRAS* reduced cell proliferation compared to the non-target (*NT*) control group (**Figure 1D**). Importantly, only *KRAS* knockdown abolished colony formation in soft agar assays (**Figures 1E-1F**). Also, after xenografting into immunocompromised mice, *NF1* KO iHSC1λ cells with *KRAS*-knockdown, but not those with *HRAS*-knockdown, showed reduced tumor-forming ability (**Figures 1G-1I**). Conversely, in WT iHSC1λ cells, *HRAS* knockdown, but not *KRAS* knockdown, ablated colony formation, although both reduced proliferation (**Figures S3G-S3I**). In summary, *NF1* KO and WT iHSC1λ cells depend mainly on KRAS and HRAS, respectively. The selective KRAS dependency in NF1-deficient SCs suggests that targeting KRAS is a potential therapeutic strategy for NF1-deficient SC tumors.

### Selective induction of pyroptosis in NF1-deficient SCs by PKC agonism

To identify therapeutics that leverage the KRAS dependency in NF1-deficient tumor cells, we focused on PKC agonists due to their preclinical therapeutic effects on KRAS mutant-driven pancreatic and colon cancers.^24, 31, 32^ Epistasis analysis of *NF1* and the nine PKC genes using The Cancer Genome Atlas (TCGA) pan-cancer dataset revealed that each PKC family gene exhibited co-mutation with *NF1* (P<0.001), implying functional dependency between PKC and NF1 in driving oncogenesis (**Figures S4A-S4C)**. Given that most cancer-associated PKC mutations are loss-of-function,^21^ we hypothesized that strategies promoting PKC activation would inhibit NF1-loss-associated tumorigenesis and tumor growth. Prostratin is a natural plant-derived PKC activator that shows anti-tumor activities in multiple types of tumors.^24, 33, 34^ We tested prostratin in iHSC1λ cells and found that prostratin selectively suppressed the viability and induced CASP3 cleavage in *NF1* KO iHSC1λ, but not in WT iHSC1λ (**Figures 2A-2B**). Furthermore, prostratin inhibited colony formation by *NF1* KO iHSC1λ, but not by WT iHSC1λ, in soft agar (**Figure 2C**). Thus, prostratin effectively and selectively inhibits growth of immortalized NF1-deficient SCs.

**Figure 2.**
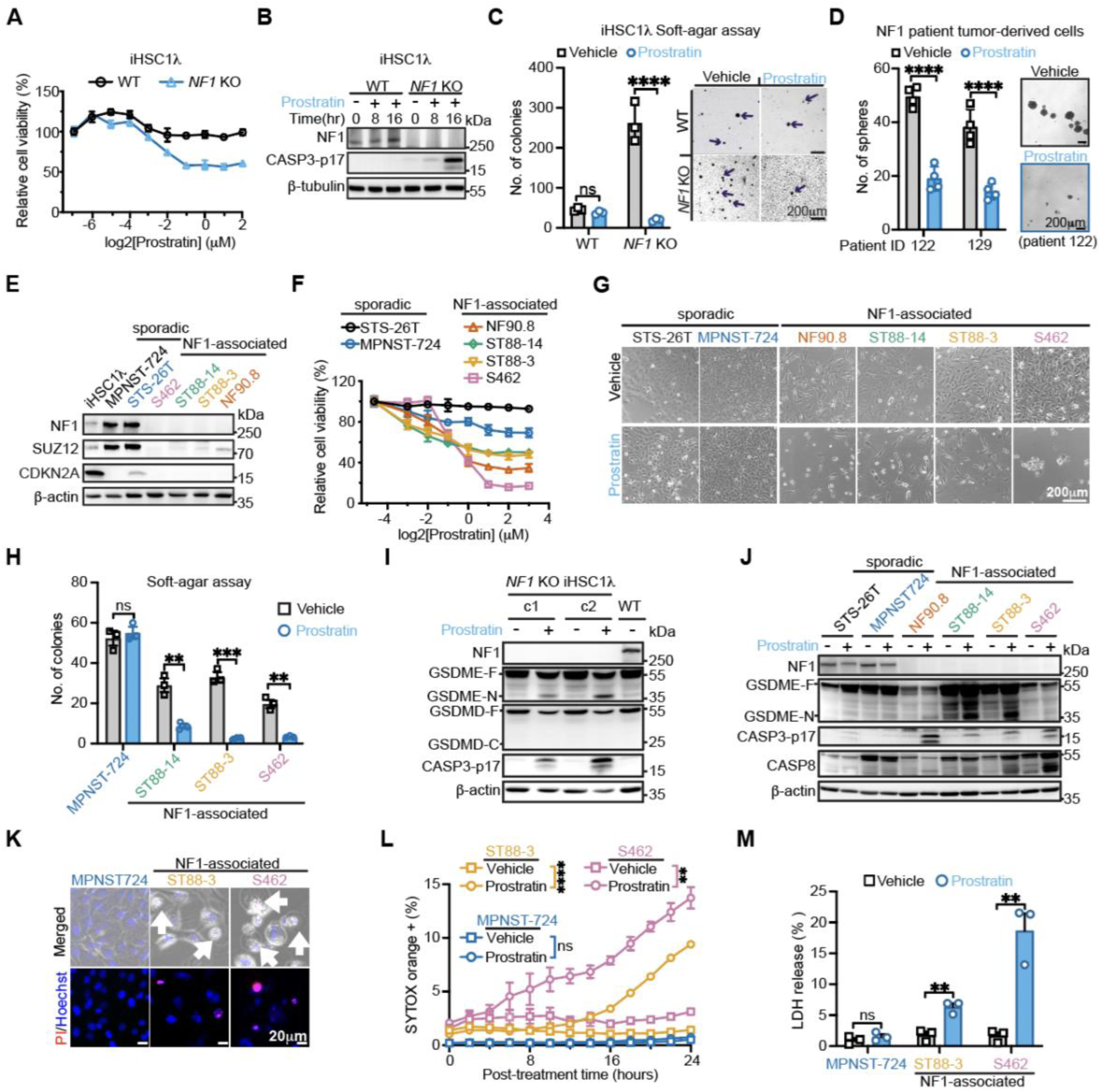
Prostratin inhibits tumorigenesis and induces pyroptosis of NF1-deficient SCs. (**A-B**) WT and *NF1* KO iHSC1λ cells: viability-based dose-response curves of prostratin (A) and immunoblotting indicated proteins following treatments (± prostratin, 1μM) for indicated hours (B). CASP3-p17: cleaved-CASP3. |(**C-D**) Soft-agar assays of iHSC1λ cells (C) and sphere formation assays in human neurofibroma samples (D) (± prostratin, 1μM). Left: quantification of colonies/spheres per well; right: representative images. Arrows indicate colonies. (**E-G**) Analysis of MPNST cell lines: Immunoblotting indicated proteins (E), prostratin dose-response curves (F), and representative phase-contrast images (± prostratin 2μM, 96 h) (G). (**H**) Soft-agar assays in MPNST cells (± prostratin, 2μM). (**I-J**) Immunoblotting indicated proteins in iHSC1λ (I) and MPNST cell lines (J) (± prostratin, 2μM). GSDME-F/GSDMD-F: full-length GSDME/GSDMD; GSDME-N: N-terminal GSDME fragment. (**K-M**) Analysis of MPNST cells: Representative images of live cells (+prostratin 2μM, 24h) with Hoechst-33342 and propidium-iodide (PI) (K), and pyroptosis assessed by SYTOX orange uptake (L) and LDH release (± prostratin 2μM, 30 h) (M). Arrows indicate bubble-like protrusions. ns, p>0.05; ∗∗p<0.01, ∗∗∗p<0.001, ∗∗∗∗p<0.0001 by t-test, except (L) by two-way ANOVA.

We next examined whether prostratin shows therapeutic effects in primary, non-immortalized, NF1-deficient tumor cells from patients. Cells derived from surgically excised human benign neurofibromas carrying *NF1* mutations were cultured as spheres and formed fewer spheres following prostratin treatment (**Figure 2D**). We next utilized six human MPNST cell lines to assess the effect of prostratin in malignant tumor cells. Four cell lines (ST88-14, ST88-3, S462, and NF90.8) derived from NF1-associated MPNST lacked NF1 expression (**Figure 2E**). Prostratin profoundly reduced viability of these NF1-associated MPNST cell lines, while causing only minor growth inhibition in the two sporadic MPNST cell lines (**Figures 2F-2G**). Additionally, prostratin significantly decreased colony numbers formed by NF1-associated MPNST cell lines, but spared colony formation in a sporadic MPNST cell line (**Figure 2H**). Together, these data demonstrate that prostratin effectively and selectively inhibits tumorigenesis-associated phenotypes in benign and malignant NF1-deficient SCs.

To determine the cell death type induced by prostratin treatment, we analyzed cleavage patterns of critical death-effector proteins (GSDME, GSDMD, and CASP3) in *NF1* KO iHSC1λ and NF1-associated MPNST cell lines. GSDME, a specific effector of pyroptosis, was cleaved in NF1-deficient cells upon prostratin treatment (**Figures 2I-2J)**. Consistent with this finding, morphological features of pyroptosis, including plasma membrane bubble-like protrusions and propidium iodide-stained nuclei, were observed in prostratin-treated NF1-deficient MPNST cells but not in sporadic MPNST cells (**Figures 2K, S5A**). Additional features of pyroptosis, including increasing SYTOX orange uptake and elevated supernatant lactate dehydrogenase (LDH) levels, were also detected in NF1-deficient cell lines following prostratin treatment, but not in a sporadic MPNST cell line (**Figures 2L-2M, S5B**). Thus, prostratin selectively induces pyroptosis in NF1-deficient SCs.

### Pyroptosis is triggered by the CASP8/CASP3/GSDME pathway

We next examined the molecular basis underlying prostratin-triggered pyroptosis in NF1-deficient SC lines. Pyroptosis is characterized by the cleavage of full-length gasdermin proteins, which can be mediated by caspases.^1, 35^ In NF1-deficient SC lines treated with prostratin, CASP3 was activated along with GSDME cleavage (**Figures 2I-2J)**. Analysis of the enzymes upstream of CASP3 (CASP8 and CASP9) in *NF1* KO iHSC1λ and the NF1-associated MPNST cell line ST88-3 showed increased CASP8 cleavage after prostratin treatment (**Figures S5C-S5D**). CASP8 and CASP3 inhibitors, but not a CASP9 inhibitor, suppressed the cleavage of CASP3 and GSDME (**Figures S5D-S5E**). These data suggest that CASP8 activation is essential for triggering pyroptosis in NF1-deficient SCs upon PKC agonism.

To confirm this, we knocked out CASP8 with CRIPSR/Cas9 in NF1-associated MPNST cell lines S462 (*C8*-KO S462) and ST88-3 (*C8*-KO ST88-3) (**Figures S5F**). In *C8*-KO S462 cells, prostratin failed to induce CASP3/GSDME cleavage or increase LDH release (**Figures S5G-S5H**), and the prostratin-mediated reduction in cell viability was rescued compared to parental S462 cells (**Figure S5I**). Similarly, *C8*-KO ST88-3 cells were protected from both prostratin-induced elevation of LDH release and inhibition of cell viability (**Figures S5J-S5K**). These data indicate that CASP8 functions upstream of CASP3/GSDME activation in prostratin-induced pyroptosis of NF1-deficient cells.

To validate this mechanism, we tested whether CASP8 activation mimics the effect of prostratin. We expressed wild-type CASP8 (CASP8-WT) or enzyme-deficient CASP8 (CASP8-C360S) (**Figure S5L**) in three NF1-deficient cell lines (*NF1* KO iHSC1λ, S462, and ST88-3). Inducible expression of CASP8-WT, but not CASP8-C360S, triggered autonomous CASP8 activation as previously reported^36^ and subsequent cleavage of CASP3 and GSDME (**Figure S5M**). CASP8-WT expression also reduced cell viability in all three SC lines, while CASP8-C360S had no effect (**Figure S5N**). Thus, CASP8 activation alone is sufficient to induce pyroptosis and impair cell viability in NF1-deficient SCs.

Next, to determine whether GSDME is required for prostratin-induced pyroptosis, we generated *GSDME* KO cell lines from NF1-deficient cells (*NF1* KO iHSC1λ, ST88-3, S462) (**Figure S6A**). Upon prostratin treatment, these *GSDME* KO lines failed to show pyroptotic features, including bubble-like membrane protrusions, propidium iodide-stained nuclei, increased SYTOX orange uptake, and elevated LDH release (**Figures S6B-S6E)**. Although pyroptosis was ablated, cell viability remained suppressed (**Figure S6F**), and the cells underwent apoptosis, exhibiting membrane blebbing, CASP8/CASP3 cleavage, and increased Annexin-V staining (**Figures S6B-S6C, S6G-S6H**). Live-cell imaging further revealed that only GSDME-expressing parental cells proceeded to pyroptosis, accounting for the majority of cell death (60-70%) upon prostratin treatment for 24h (**Figures S6I-S6M**). Thus, prostratin triggers the CASP8/CASP3/GSDME pathway to induce pyroptosis in NF1-deficient SCs, with GSDME expression determining the pyroptotic fate.

### Induction of pyroptosis in NF1-deficient SCs by PKCδ

Prostratin is a broad-spectrum PKC agonist.^24^ We sought to determine the PKC isozyme(s) that induce pyroptosis in SCs with NF1 loss. Of the nine PKC members, *PKC*α, δ, ε, η, and ι mRNAs were expressed in all eight cell lines tested (**Figure 3A**). To identify relevant isozymes, we combined prostratin with PKC inhibitors, either Go6976 (inhibitor of PKCα/β/γ) or Go6983 (inhibitor of PKCα/β/γ and PKCδ/ε/η/θ).^37^ Results from NF1-deficient cells showed that only the broad-spectrum inhibitor Go6983 rescued the cell viability inhibitory effect of prostratin (**Figure S7A**) and abrogated pyroptosis induction (**Figure 3B**). Given the PKC expression profile in SC lines and the inhibitory profiles of Go6976 and Go6983, these data suggest that one or more of the novel PKC isozymes (δ, ε, η) mediates pyroptosis in NF1-deficient SCs.

**Figure 3.**
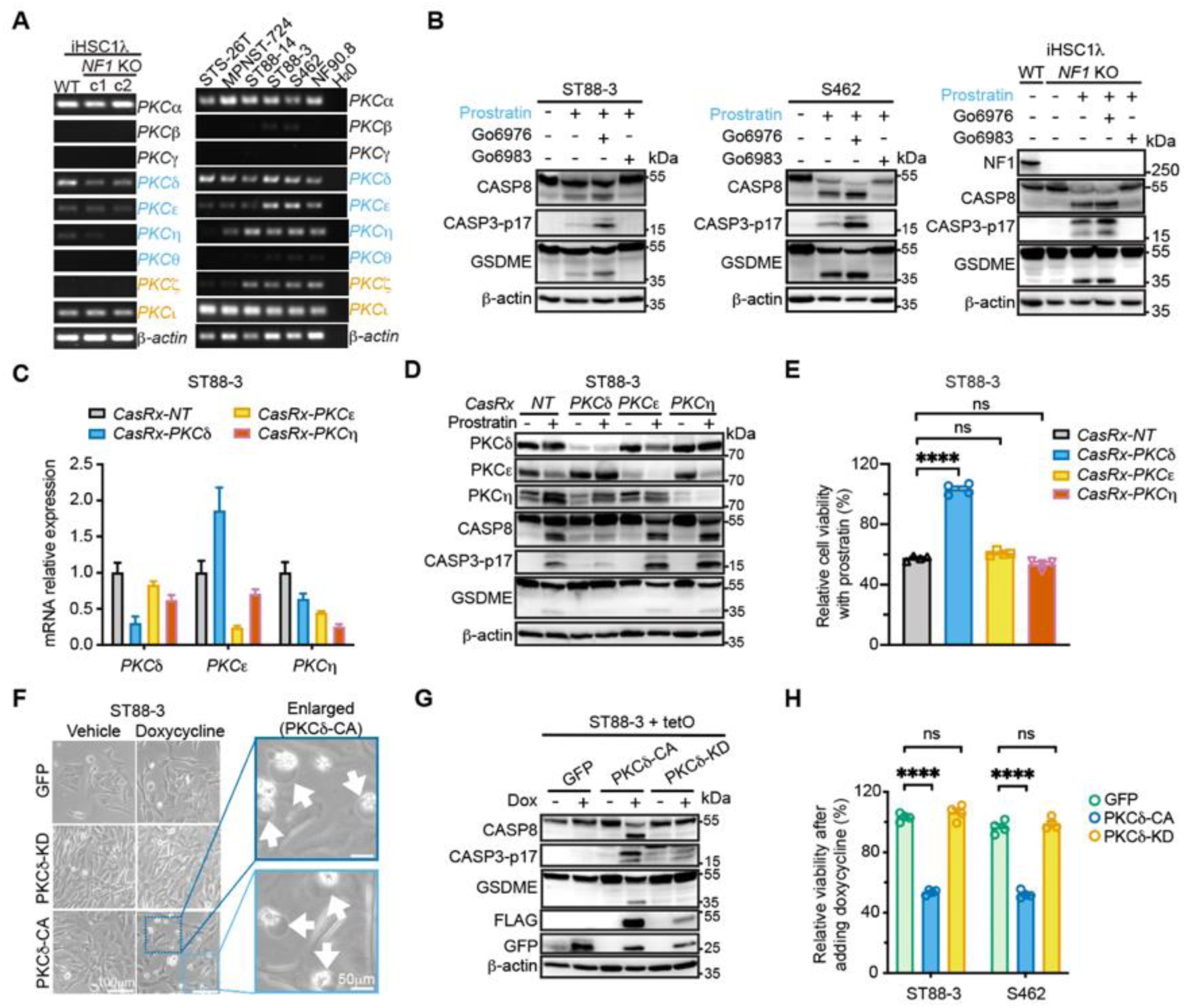
PKCδ mediates prostratin-induced pyroptosis in NF1-deficient SCs. (**A**) RT-PCR analysis of PKC isozyme expression in iHSC1λ and MPNST cell lines. (**B**) Immunoblotting indicated proteins in cell lines treated with vehicle, prostratin (2μM), or prostratin plus PKC inhibitors Go6976 (2μM) or Go6983 (1μM). (**C-E**) *PKCδ/ε/η* knockdown in ST88-3 cells: qPCR analysis of mRNA levels (C), immunoblotting indicated proteins (± prostratin 2μM) (D), and relative cell viability of cell lines with individual *PKCδ/ε/η* knockdown (± prostratin 2μM, 96h), normalized to vehicle (E). NT: non-target. (**F-G**) ST88-3 cells with inducible expression of PKCδ-mutant or GFP treated with vehicle or doxycycline (1μg/ml): representative images (F) and immunoblotting indicated proteins (G). CA: constitutive-active; KD: kinase-dead (PKCδ-K378R). Arrows indicate air-bubble structures. (**H**) Relative viability of ST88-3 and S462 cells with inducible expression of PKCδ-mutant or GFP following doxycycline treatment (1μg/ml, 72h). ns, p>0.05; ∗∗∗∗p<0.0001 by ANOVA.

We next employed the CasRx system to knock down *PKCδ*, *PKCε*, or *PKCη* in NF1-deficient cell lines (ST88-3 and S462) (**Figures 3C, S7B**). Notably, the knock-down of *PKCδ*, but not *PKCε* or *PKCη*, reduced prostratin-induced cleavage of CASP8/CASP3/GSDME (**Figures 3D, S7C**). *PKCδ* knockdown also rescued the prostratin-induced cell viability inhibition (**Figures 3E, S7D).** These data suggest that PKCδ mediates prostratin-induced pyroptosis in NF1-deficient SCs. Indeed, a modest increase in PKCδ phosphorylation at S664, which suggested PKCδ activation, was observed in prostratin-treated NF1-deficient SCs (**Figures S7E-S7F**). We next tested whether PKCδ activation recapitulated the effects of prostratin. Expression of constitutively active PKCδ (PKCδ-CA), but not kinase-dead PKCδ (PKCδ-KD)^38^ or GFP control, in NF1-deficient MPNST cell lines led to the formation of bubble-like protrusions, cleavage of CASP8/CASP3/GSDME, and inhibition of cell viability (**Figures 3F-3H**). Similarly, the expression of PKCδ-CA, but not PKCδ-KD or GFP, in *NF1* KO iHSC1λ cells activated CASP3 and impaired cell viability (**Figures S7G-S7H**). Taken together, these data indicate that PKCδ agonism induces pyroptosis in NF1-deficient SCs.

### Pyroptosis requires KRAS phosphorylation

Given that PKCδ kinase activity is required to induce pyroptosis in NF1-deficient SCs, we sought to identify its relevant substrate(s). We hypothesized that KRAS is a PKCδ substrate mediating pyroptosis because KRAS is a known direct substrate of conventional PKCs^39^, and NF1-deficient SCs are KRAS-dependent (**Figure 1**). Supporting this, the knock-down of *KRAS,* but not *HRAS* or *NRAS,* rescued prostratin-induced cell viability inhibition in *NF1* KO iHSC1λ cells (**Figure 4A**). Furthermore, prostratin treatment induced colocalization of PKCδ with KRAS, but not with red fluorescence protein (RFP) or HRAS, at plasma membrane and perinuclear structures (**Figures 4B, S8A-S8E**), suggesting a functional connection between PKCδ and KRAS.

**Figure 4.**
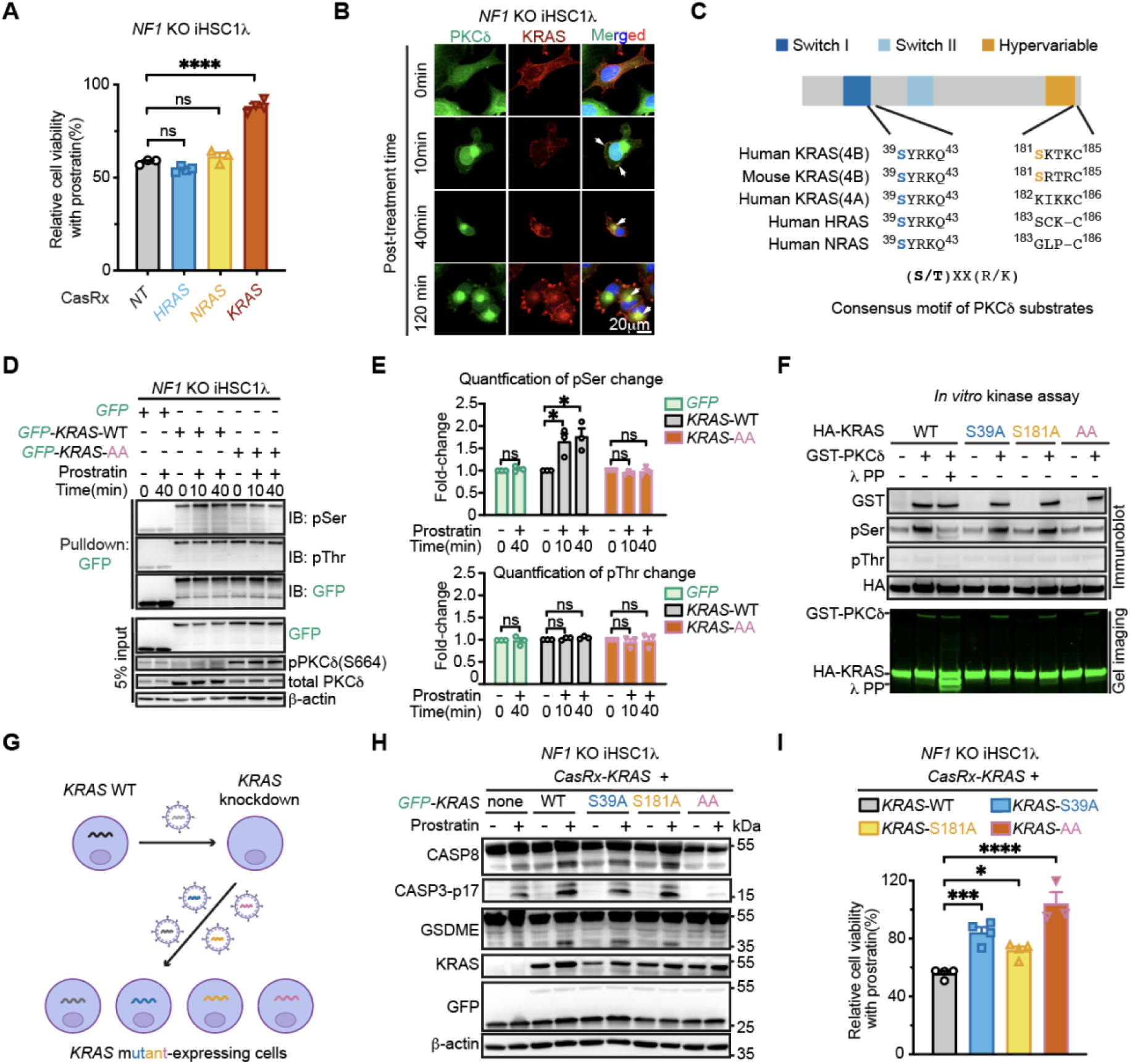
KRAS phosphorylation is essential for prostratin-induced pyroptosis in NF1-deficient SCs. **(A)** Relative cell viability of *NF1* KO iHSC1λ with individual knockdown of *H-/N-/K-RAS* or NT (± prostratin, 2μM), normalized to vehicle. **(B)** Representative images of PKCδ-GFP and mRuby2-KRAS4B in *NF1* KO iHSC1λ cells (± prostratin, 2μM). Arrows indicate colocalization. **(C)** PKCδ phosphorylation site analysis in RAS paralogs. (**D-E**) Immunoblotting serine/threonine phosphorylation in GFP-immunoprecipitations from indicated iHSC1λ cell lines (D) and the quantifications (E). Cells were starved and then stimulated for indicated minutes (±prostratin 2.5μM). AA: S39A/S181A. (**F**) Immunoblotting and direct gel imaging of PKCδ, λ protein phosphatase (λPP), HA-tagged KRAS4B mutants after *in vitro* kinase assays (n=2). (**G-I**) KRAS-WT replacement with KRAS-mutant in *NF1* KO iHSC1λ cells: the “knock-down and add-back” strategy (G), immunoblotting indicated proteins (H), and relative cell viability (± prostratin, 2.5μM) (I). ns, p>0.05; ∗p < 0.05, ∗∗∗p<0.001, ∗∗∗∗p<0.0001 by ANOVA.

To identify potential PKCδ phosphorylation sites in KRAS, we analyzed the PKCδ substrate consensus motif (S/TXXR/K)^40^ in KRAS4B, the only KRAS isoform expressed in SCs (**Figure S8F**). Two serine residues (S39 and S181) were predicted as potential phosphorylation sites with high probability scores (>0.90) (**Figures 4C, S8G**). Notably, KRAS-S181 phosphorylation by conventional PKC was previously detected in fibroblasts.^39^ Phosphoproteomics analysis confirmed that prostratin induced KRAS-S39 phosphorylation (**Figure S8H**); S181 phosphorylation was not detected by this approach for technical reasons detailed in the Methods. Consistent with these findings, prostratin treatment increased total serine, but not threonine, phosphorylation of KRAS in NF1-deficient cells (**Figures 4D-4E, S8I-S8J**). To confirm that both sites contribute to KRAS phosphorylation, we generated a double-non-phosphorylatable KRAS-S39A/S181A (KRAS-AA) mutant. The increased serine phosphorylation was abolished in this mutant and was absent in HRAS (**Figures 4D-4E, S8I-S8L**). However, prostratin-induced phosphorylation remained detectable in single-non-phosphorylatable KRAS-S39A and KRAS-S181A mutants (**Figure S8M**). Corroborating these findings, *in vitro* kinase assays showed that PKCδ increased total serine phosphorylation of KRAS-S39A and KRAS-S181A mutants, but not the KRAS-AA mutant (**Figure 4F**). Thus, PKCδ directly phosphorylates KRAS4B at both S39 and S181 residues.

To characterize the functional significance of these two KRAS phosphorylation sites, we employed a “knock-down and add-back” strategy, substituting wild-type KRAS (KRAS-WT) with KRAS phospho-mutants (**Figure 4G**). While individual KRAS-S39A or KRAS-S181A mutants showed minimal effects, the double-non-phosphorylatable mutant KRAS-AA abolished prostratin-induced cleavage of CASP8/CASP3/GSDME (**Figure 4H**). Consistent with these findings, KRAS-AA substantially rescued cell viability in prostratin-treated *NF1* KO iHSC1λ cells (**Figure 4I**). Similar protective effects of KRAS-AA on pyroptosis and cell viability were observed in ST88-3 and S462 cells (**Figures S8K-S8M**). Taken together, KRAS phosphorylation at S39 and S181 by PKCδ is necessary for prostratin-induced pyroptosis in NF1-deficient SCs.

### Dual phosphorylation promotes KRAS-GDP accumulation and KRAS-CASP8 interaction

Previous studies showed that KRAS-S181 phosphorylation could cause KRAS translocation to the endoplasmic reticulum (ER), and that ER-localized KRAS with S181 phosphorylation triggers cell death in fibroblasts and colon cancer cells.^31^ Consistent with this, in *NF1* KO iHSC1λ and S462 cells, prostratin treatment induced robust translocation of KRAS-WT to the ER (**Figures S9A-S9B**), colocalizing with BAP31 and calnexin while showing minimal colocalization with the mitochondrial marker TOMM20 by super-resolution microscopy (**Figures S9C-S9E**). By contrast, the phosphorylation-deficient KRAS-AA mutant exhibited minimal ER localization following prostratin treatment (**Figures S9F-S9G**). The dual-phosphomimetic KRAS-S39E/S181E (KRAS-EE) mutant was constitutively localized to the ER but not mitochondria, independent of prostratin treatment (**Figures S9H-S9I**). Further analysis revealed that KRAS-S39E and KRAS-S181E each promoted ER localization, with KRAS-S181E showing a stronger effect (**Figure S9J**). These data indicate that dual phosphorylation promotes KRAS translocation to the ER, with S181 as the primary determinant, consistent with previous findings.^24, 32^ Notably, NF1-deficient cells expressing KRAS-EE or KRAS-S181E remained viable and proliferative in the absence of prostratin treatment (**Figures S9H-S9J**), demonstrating that ER translocation of KRAS is necessary but not sufficient for cell death induction, and that additional prostratin-induced molecular events are required—distinct from prior observations in which KRAS-S181E alone trigger autophagic cell death in fibroblasts and colon cancer cells^31^ as detailed in Discussion.

To investigate the molecular basis by which dually phosphorylated KRAS promotes cell death, we first excluded ER stress as an alternative mechanism, as prostratin treatment did not activate unfolded protein response (UPR) markers CHOP, XBP1s, or ATF4 in NF1-deficient cells (**Figure S9K**). Given that CASP8 translocates to the ER in response to cell death-inducing stimuli,^41^ we investigated a potential KRAS-CASP8 interaction. CASP8 localized to both the ER and mitochondria in NF1-deficient cells (**Figure S9L**), whereas phosphorylated KRAS localized selectively to the ER but not mitochondria (**Figures S9C-S9E, S9J**). Since KRAS’s ER-restricted localization overlaps with only a subset of CASP8’s subcellular distribution, KRAS phosphorylation thus determines the specific subcellular compartment (the ER) at which KRAS–CASP8 complex formation occurs. Consistent with this, prostratin induced colocalization of KRAS-WT, but not the phosphorylation-deficient KRAS-AA mutant, with CASP8 at the ER (**Figures 5A, S9M–S9O**). Furthermore, prostratin enhanced KRAS-WT interaction with CASP8 in NF1-deficient SC lines, while KRAS-AA showed no such enhancement (**Figures 5B, S9P**). Unexpectedly, CASP8 preferentially interacted with GDP-bound KRAS-WT (KRAS-WT-GDP) over GTP-bound KRAS-WT (KRAS-WT-GTP), contrasting with the canonical RAS effector RAF1 that preferentially binds KRAS-GTP (**Figure 5C**). Supporting this GDP preference, prostratin treatment reduced KRAS-GTP levels in NF1-deficient cells (**Figure 5D**). Importantly, the phosphomimetic KRAS-EE exhibited stronger CASP8 interaction than KRAS-WT, particularly when GDP-bound, recapitulating the effect of prostratin (**Figures 5E, S9Q**). While prostratin enhanced KRAS-WT/CASP8 interaction in both nucleotide-bound states, it had no effect on KRAS-AA/CASP8 interaction (**Figures 5B, S9R**). Together, these data demonstrate that prostratin-induced dual phosphorylation promotes KRAS-GDP interaction with CASP8 at the ER, and that this interaction is the critical event linking KRAS phosphorylation to cell death.

**Figure 5.**
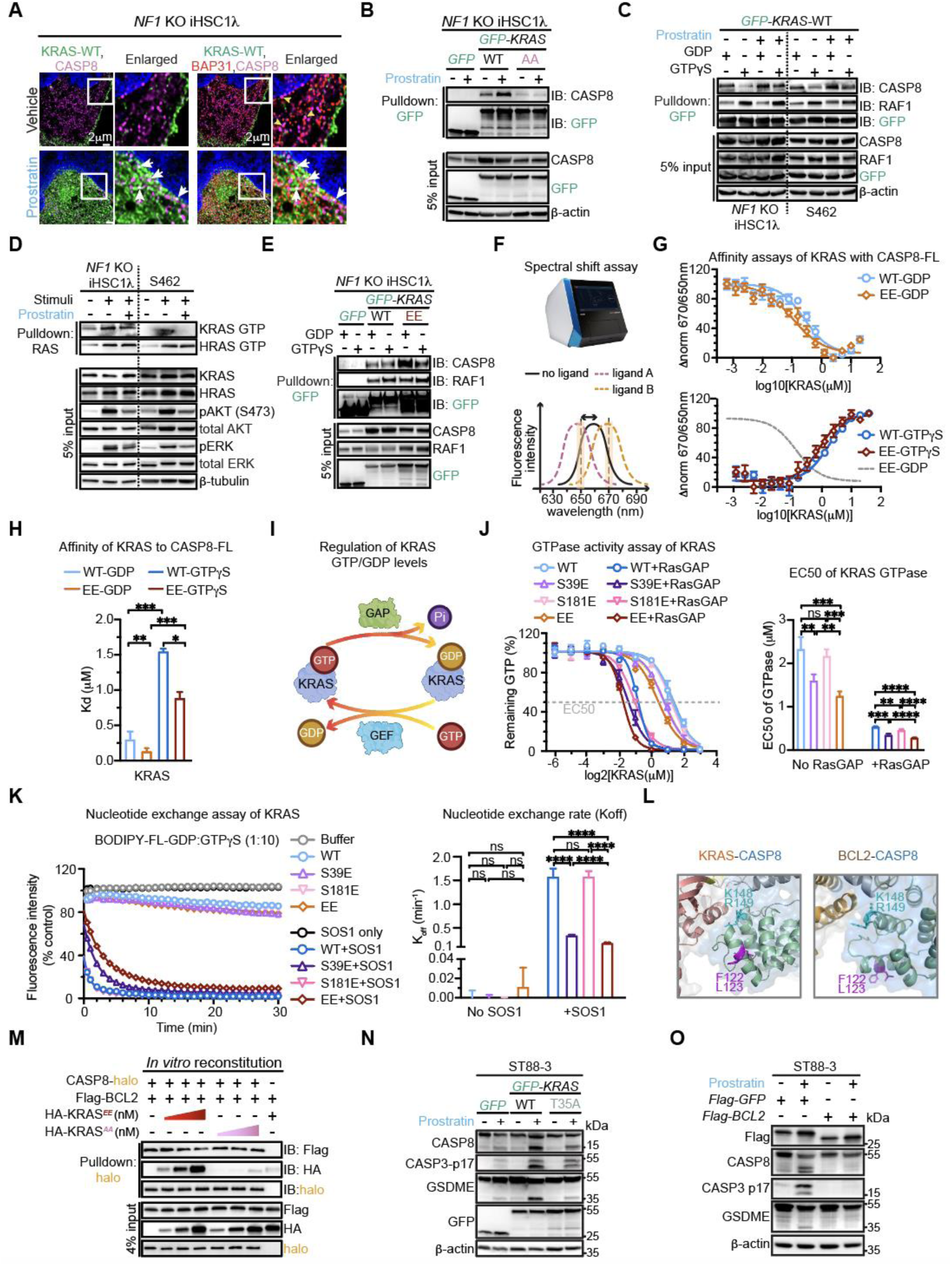
Phosphorylation modulates KRAS localization and KRAS-GDP levels, promoting CASP8 interaction and activation. (**A**) Representative super-resolution microscopic images of GFP-KRAS-WT, CASP8-C360S, and BAP31 in *NF1* KO iHSC1λ (± prostratin 2μM, 13 h). Arrows (white) indicate KRAS/CASP8 and KRAS/BAP31/CASP8 colocalization; arrowheads (yellow) show CASP8/BAP31 colocalization. (**B-C**) Immunoblotting of GFP-immunoprecipitations from indicated cell lines (±prostratin 2μM, 13 h) (B) and cellular lysates subjected to GDP/GTPγS loading (±prostratin 2μM) (C). **(D)** RAS pull-down assays following serum stimulation (15%, 5min) in pre-treated cells (± prostratin 2.5μM, 13h). **(E)** Immunoblotting of GFP-immunoprecipitations from lysates of indicated cell lines subjected to GDP/GTPγS loading. (**F-H**) Affinity analysis of KRAS variants with RED-NHS-labeled full-length CASP8 (CASP8-FL): schematic spectral-shift assay (F), KRAS-CASP8 affinity assays (n=3-4) (G) and Kd comparisons (H). (**I-K**) RAS variant characterization: Schematic RAS-GDP/-GTP regulation (I), GTPase hydrolysis assays with EC_50_ quantification (n=3) (J), and nucleotide exchange assay with K_off_ quantification (n=2) (K). (**L**) Interface analysis of predicted KRAS-CASP8 and BCL2-CASP8 complexes. (**M**) Immunoblotting analysis of competition assays. CASP8-FL-halo/Flag-BCL2 complexes were titrated with HA-KRAS-EE-GDP or HA-KRAS-AA-GTPγS. (**N-O**) Immunoblotting indicated proteins in cells expressing GFP, GFP-KRAS-WT/-T35A (N), or Flag-GFP/-BCL2 (O) (±prostratin 2μM, 24h). ns, p>0.05; ∗∗p<0.01, ∗∗∗p<0.001, ∗∗∗∗p<0.0001 by ANOVA.

To validate these findings, we performed *in vitro* protein binding assays using purified recombinant KRAS and CASP8 proteins. Mammalian-derived recombinant KRAS-WT and KRAS-EE proteins were validated for nucleotide-bound states using RAS pulldowns and microscale thermophoresis affinity assays, confirming that GTP-bound KRAS proteins exhibit higher affinity for RAF1 RAS-binding domain (RAF1-RBD) than GDP-bound forms (**Figures S10A-S10C**). Due to homo-oligomerization and self-cleavage of CASP8-WT,^42^ we engineered an oligomerization/cleavage-deficient CASP8-full-length (CASP8-FL) mutant carrying F122G/L123G/D374A/D384A mutations (**Figure S5J**)^42, 43^ and purified it from a cell-free protein expression system (**Figures S10D-S10F**). Spectral-shift assays^44^ with purified proteins revealed that KRAS-EE displayed higher affinity for CASP8-FL than for KRAS-WT in either nucleotide state, with GDP-bound KRAS-EE showing maximal affinity (**Figures 5F-5H**). Collectively, these results demonstrate that dually phosphorylated KRAS-GDP interacts with CASP8, establishing CASP8 as a functional KRAS-GDP effector.

We next investigated whether KRAS phosphorylation directly modulates KRAS-GTP levels, which are regulated through two mechanisms: GTP hydrolysis and GDP dissociation (**Figure 5I**).^18^ KRAS-EE exhibited the highest rate of intrinsic GTP hydrolysis, followed by KRAS-S39E, while KRAS-S181E showed rates comparable to KRAS-WT (**Figure 5J**). Under RasGAP (p120GAP)-stimulated conditions, a similar pattern emerged, except that KRAS-S181E also displayed enhanced hydrolysis relative to KRAS-WT (**Figure 5J**). Assessment of GDP dissociation kinetics showed that all KRAS variants had similar intrinsic rates (**Figure 5K**); however, in the presence of the RAS guanine exchange factor (RasGEF) SOS1, KRAS-EE and KRAS-S39E—but not KRAS-S181E—showed reduced GDP dissociation compared to KRAS-WT (**Figure 5K**). These findings suggest that KRAS phosphorylation, particularly at S39, promotes KRAS-GDP accumulation via two complementary mechanisms: enhanced GTPase activity and impaired RasGEF-induced nucleotide exchange, thereby facilitating KRAS-GDP/CASP8 interaction.

To understand the structural basis of KRAS-CASP8 interaction, we employed AlphaFold-2 to predict this complex structure (**Figure S11A**). The predicted complex suggested engagement of both CASP8 N-terminal (Nter, DED-DED domains) and C-terminal (Cter, p18-p10 domains) lobes with KRAS (**Figures S11B-S11C**). Cellular KRAS pulldown assays confirmed this prediction, showing that KRAS-EE interacted with both CASP8 truncation mutants, albeit with lower affinity compared to CASP8-FL (**Figures S11D-S11F**). Affinity assays with purified CASP8 truncation mutants further confirmed that both lobes interacted with GDP-bound KRAS-EE, with CASP8-Nter exhibiting stronger affinity (**Figures S11G-S11J**). CASP8-K148/R149, located within the CASP8-Nter, were predicted as interaction interfaces (**Figure S11K**). Notably, these residues also mediate CASP8 oligomerization (**Figure S11E**), and their mutation to K148D/R149E abolished the CASP8-Nter/KRAS-EE interaction (**Figure S11L**). Moreover, KRAS-T35, within the Switch-I domain (residues 30-40) critical for RAF1 binding,^18^ participated in the KRAS-CASP8-Cter interface (**Figure S11M**), and the KRAS-T35A mutation impaired KRAS interactions with both CASP8-Cter and CASP8-FL (**Figures S11N-S11O**). Together, these analyses define the structural and molecular basis of KRAS-CASP8 interaction.

### KRAS and BCL2 compete for CASP8 binding to control CASP8 activation

We next explored how KRAS-GDP regulates CASP8 activation. CASP8 activation has been reported to be negatively regulated by BCL2 family proteins, which localize to the ER^45^ and interact with CASP8.^46–48^ Among the three major BCL2 family members (BCL2, Bcl-xL, and MCL1), only BCL2 bound to CASP8 in NF1-deficient cells (**Figure S12A**). BCL2 predominantly localized to the ER in these cells, with colocalization with ER markers BAP31 and Calnexin, while showing minimal colocalization with the mitochondrial marker TOMM20 (**Figures S12B-S12C**), consistent with predominant ER localization of BCL2 reported in certain cell types.^49, 50^ AlphaFold prediction of the BCL2-CASP8 complex suggested an interaction between BCL2 and CASP8 (**Figure S12D**), and affinity assays confirmed that CASP8 indeed interacted with BCL2 (**Figure S12E**). Structural analysis of the predicted complex identified BCL2-R109/R110 and CASP8-K148/R149 as key interface residues (**Figure S12D**), which we validated by mutagenesis. BCL2-R109A/R110A mutations abolished BCL2 interaction with both CASP8-FL and CASP8-Nter, while BCL2 BH3-domain deletion enhanced this interaction (**Figures S12F-S12G**), demonstrating that BCL2 engages CASP8 through a non-canonical R109/R110 surface rather than its BH3-binding groove. Moreover, CASP8-K148D/R149E mutations impaired CASP8-FL/BCL2 interactions (**Figures S12H-S12I**), consistent with the AlphaFold-predicted interface (**Figure S12D**). Notably, this same CASP8-K148/R149 surface also mediates the KRAS-CASP8 complex (**Figure 5L**). Together, these data provide the spatial and structural basis for potential competition between BCL2 and KRAS for CASP8 binding.

Supporting this competition model, prostratin reduced the BCL2-CASP8 interaction while enhancing KRAS-CASP8 binding in CASP8 pulldown assays (**Figure S12A**). Importantly, *in vitro* competition assays using purified proteins (**Figures S10H-S10I**) showed that GDP-bound KRAS-EE, but not GTP-bound KRAS-AA, displaced BCL2 from CASP8 and increased its own binding to CASP8 (**Figure 5M**). Functionally, KRAS-WT overexpression enhanced prostratin-induced CASP8/CASP3/GSDME activation and cell viability suppression, while the CASP8-binding-defective KRAS-T35A mutant showed minimal effects (**Figures 5N, S12J-S12K**). Moreover, cells overexpressing KRAS-EE exhibited stronger cell death and viability reduction following prostratin treatment, compared to those expressing KRAS-WT (**Figures S12L-S12M**). Conversely, BCL2 overexpression prevented CASP8/CASP3/GSDME activation and partially rescued cell viability following prostratin treatment (**Figures 5O, S12N-S12O**). The BCL2-specific inhibitor venetoclax, disrupting the BCL2-CASP8 interaction (**Figure S12P**), enhanced prostratin-induced CASP8/CASP3/GSDME activation and viability inhibition (**Figures S12Q-S12R**). These findings demonstrate that KRAS phosphorylation promotes CASP8 activation, which involves competitive displacement of inhibitory BCL2 by dually phosphorylated KRAS. Given that dually phosphorylated KRAS-GDP preferentially binds CASP8 compared to KRAS-GTP (**Figures 5E-5H**), these results establish CASP8 as a functional KRAS-GDP effector.

### Conservation of pyroptotic vulnerability across NF1-deficient cancers

To assess if PKC agonism is effective in cancer types with prevalent *NF1* mutations, we first studied breast cancers. Neurofibromatosis type 1 patients are significantly predisposed to breast cancer,^8, 9^ and somatic *NF1* mutations are identified in 3%-18% of sporadic breast cancers, particularly in metastatic cases.^17, 51, 52^ Of the three human breast cancer cell lines tested, the triple-negative breast cancer cell line MDA-MB-231 showed absent NF1 expression (**Figure 6A**) and was more sensitive to prostratin-mediated cell viability inhibition compared to NF1-expressing lines (**Figure 6B**). Morphologically, prostratin selectively induced balloon-like membrane protrusions of MDA-MB-231 cells, suggesting pyroptosis (**Figure 6C**). As observed in SCs, prostratin induced cleavage of CASP8/CASP3/GSDME, effects enhanced by venetoclax (**Figure 6D**). The combination of venetoclax with prostratin caused stronger inhibition of cell viability in MDA-MB-231 (**Figure 6E**). Moreover, the activation of the CASP8/CASP3 axis was essential for GSDME cleavage in NF1-deficient MDA-MB-231 cells (**Figure 6F**). Knocking out *GSDME* in MDA-MB-231 prevented the formation of balloon-like membrane protrusions and suppressed the LDH release upon prostratin treatment (**Figures S13A-S13C**), suggesting GSMDE-dependent pyroptosis induced by prostratin. Notably, in *GSDME* KO MDA-MB-231 cells, prostratin induced apoptosis, as indicated by membrane blebbing, CASP8/CASP3 cleavage, and viability inhibition (**Figures S13B, S13D-S13E**). Furthermore, prostratin selectively reduced KRAS-GTP levels and increased the association of KRAS with CASP8, concomitant with decreasing interaction of BCL2 (**Figures 6G, S13F**). Importantly, prostratin, especially when combined with venetoclax, reduced the number of MDA-MB-231 colonies in soft-agar assays (**Figure S13G**). Together, these findings demonstrate that prostratin selectively triggers GSDME-dependent pyroptosis in NF1-deficient breast cancer cells and effectively inhibits tumor-relevant phenotypes.

**Figure 6.**
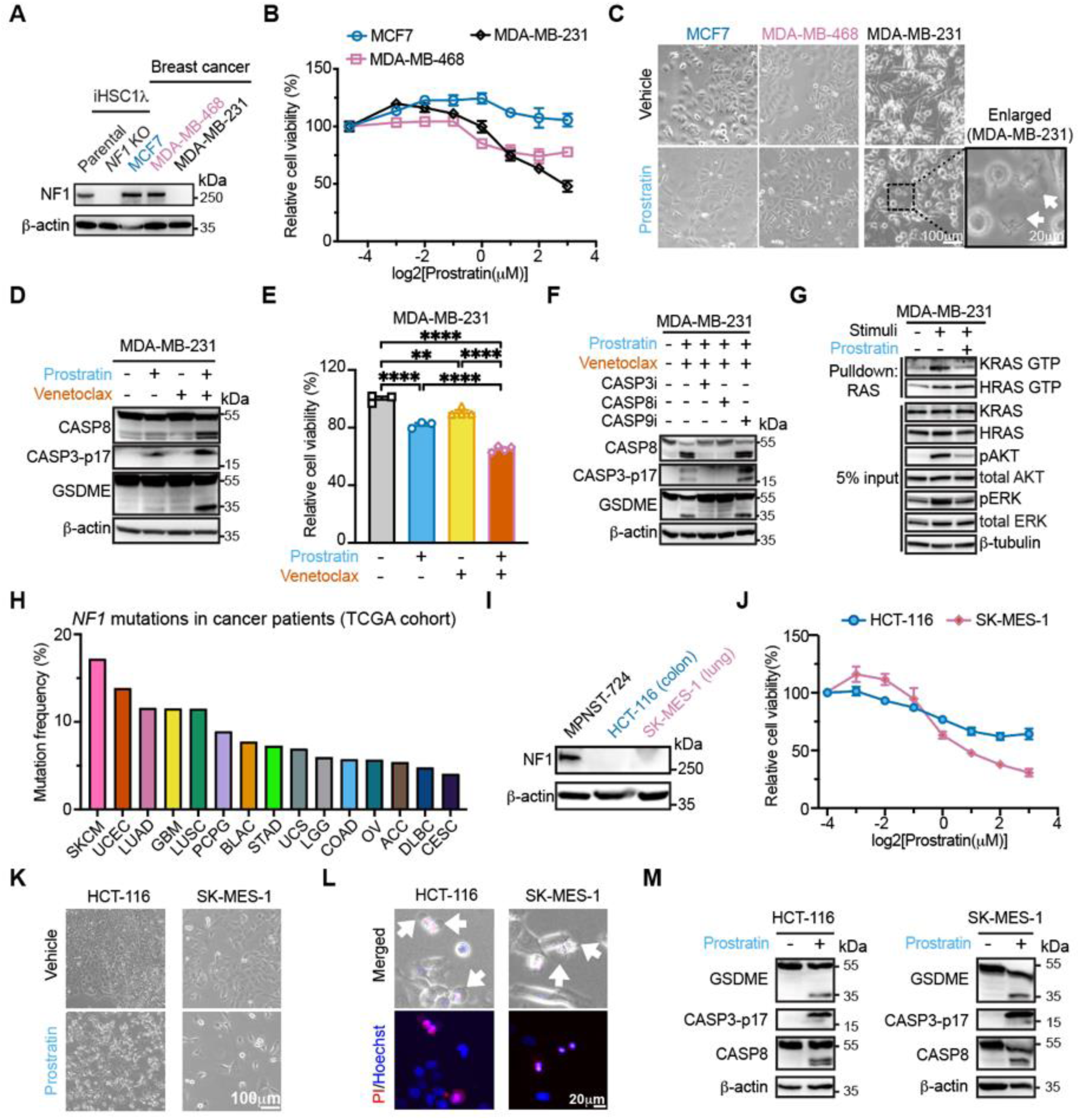
PKC agonist induces pyroptosis in multiple tumor types with NF1 deficiency. (**A-B**) Immunoblotting analysis (A) and prostratin dose-response curves (B) in indicated cell lines. (**C**) Representative images of indicated cell lines (± prostratin 4μM, 72h). (**D-E**) Immunoblotting analysis (D) and relative viabilities (E) of MDA-MB-231 upon indicated treatments (prostratin 4μM ± venetoclax 2μM), normalized to vehicle. (**F-G**) Immunoblotting analysis (F) and RAS pull-down assays (G) of MDA-MB-231 cells with indicated treatments. (**H**) Top 15 cancer types with frequent *NF1* mutations in TCGA Dataset. See **Table S1** for abbreviations. (**I-J**) Immunoblotting analysis (I) and prostratin dose-response curves (J) in indicated cell lines. (**K**) Representative phase-contrast images of indicated cell lines (±prostratin 4μM, 96h). (**L**) Representative images of cells with Hoechst-33342/PI (+prostratin 4μM, 36h). Arrows indicate bubble-like protrusions. (**M**) Immunoblotting analysis in indicated cell lines (±prostratin 4μM). ∗∗p<0.01, ∗∗∗∗p<0.0001 by one-way ANOVA.

We extended our findings to additional cancer types with frequent sporadic *NF1* mutations. Both lung and colon cancers show a relatively high frequency of *NF1* mutations (∼12% and ∼6%, respectively) in the TCGA dataset (**Figure 6H**). Two cancer cell lines, HCT-116 (colorectal adenocarcinoma) and SK-MES-1 (squamous cell lung cancer), showed NF1 loss (**Figure 6I**). Both cell lines displayed decreased viability upon prostratin treatment (**Figures 6J-6K**). Importantly, prostratin induced pyroptosis in HCT-116 and SK-MES-1 cells, as evidenced by large bubble-like membrane protrusions and GSDME cleavage (**Figures 6L-6M**). To summarize, these data demonstrate the conservation of cell death program via PKC agonism across diverse cell types with NF1 loss.

### PKC agonism inhibits NF1-deficient tumor growth *in vivo*

Mouse *Nf1* KO primary SCs exhibited similar patterns as human *NF1* KO SCs, showing increased activation of HRAS and KRAS, and PKC agonism-induced growth inhibition (**Figures S14A-S14C**). To evaluate the PKC agonism efficacy *in vivo*, we first utilized a mouse model of paraspinal plexiform neurofibroma, in which *Nf1* is deleted in *Dhh*-expressing SCs (*Dhh^Cre^;Nf1^fl/fl^* mice).^53^ Short-term prostratin treatment induced pronounced CASP3 cleavage and co-labeling of cleaved CASP3 with the SC marker CNPase in the plexiform neurofibroma tissue, indicating the death of tumorigenic SCs (**Figures S14D-S14F**). Two months of prostratin administration reduced tumor number and tumor size (**Figures 7A-7D**). Immunostaining of tumor sections revealed a decreased percentage of proliferative Ki-67-positive cells and increased staining of cleaved CASP3 following prostratin treatment (**Figures 7E-7F**).

**Figure 7.**
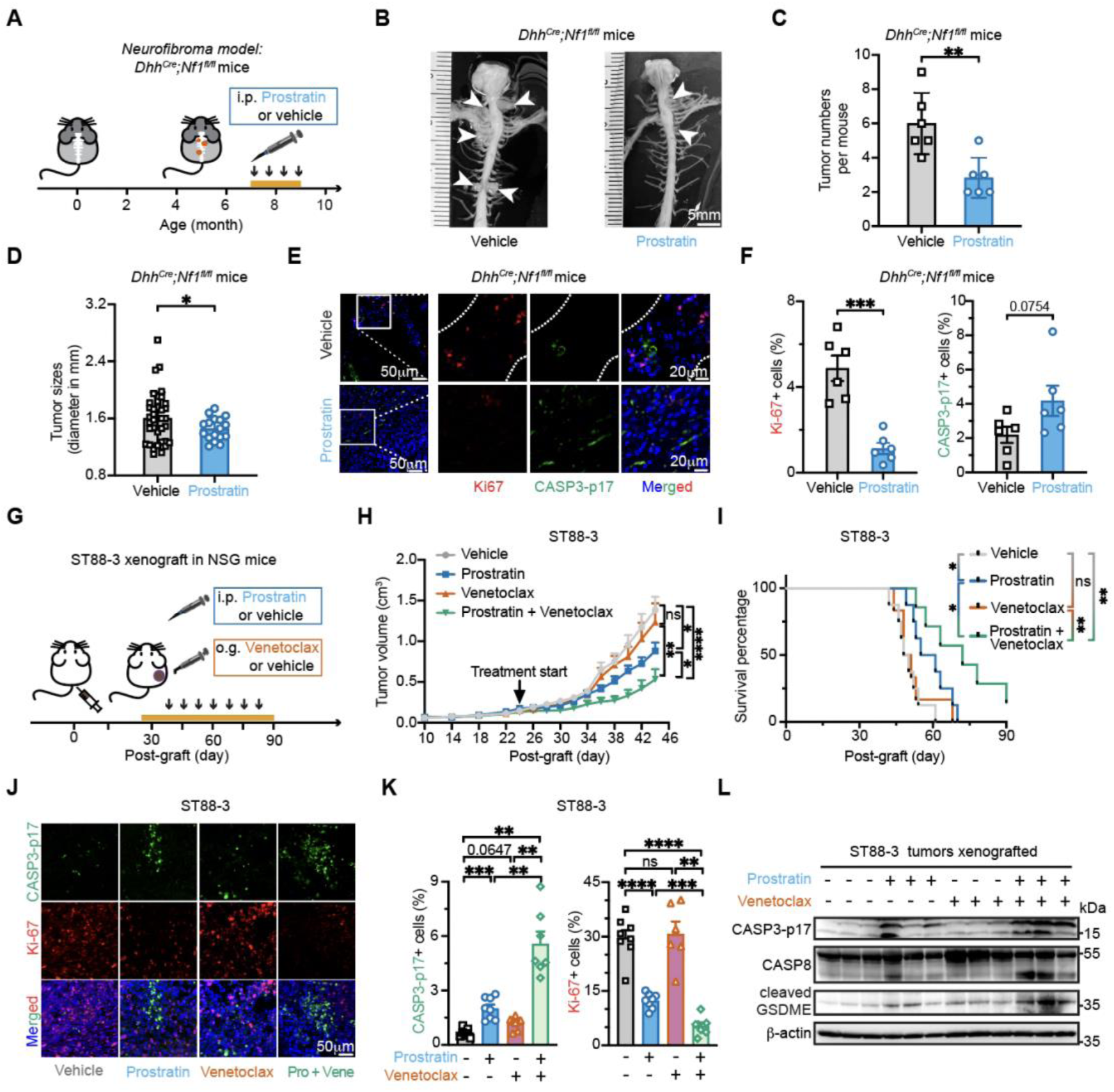
PKC agonist exhibits therapeutic efficacy in NF1-deficient tumors *in vivo*. (**A**) Schematic two-month treatment (yellow bar) with vehicle or prostratin (0.8mg/kg) in *Dhh^Cre^;Nf1^fl/fl^* mice. (**B-D**) Representative images of plexiform neurofibroma (B) with quantification of tumor numbers (C) and sizes (D) from treated mice in (A) (n=6/group). Arrowheads indicate neurofibromas. (**E-F**) Representative immunostaining images of Ki-67 and CASP3-p17 (E) with quantification (F) in treated-neurofibromas. (**G**) Experimental design for panels (H-L). (**H-I**) Tumor growth kinetics (H) and survival analysis (I) of ST88-3 grafted NSG mice treated with vehicle (n=8), prostratin (n=8), venetoclax (n=6), or Prostratin+Venetoclax (n=7). (**J-L**) Representative immunostaining images of CASP3-p17 and Ki-67 (J) with quantification (K), and immunoblotting analysis (L) of treated ST88-3 tumors. ns, p>0.05; ∗p<0.05, ∗∗p<0.01, ∗∗∗p<0.001, ∗∗∗∗p<0.0001 by t-test (C-F), two-way ANOVA(H), log-rank test(I), and one-way ANOVA(K).

Next, we studied the efficacy of prostratin in human MPNST xenografts (**Figure 7G**). In NF1-deficient ST88-3 xenografts, prostratin suppressed tumor growth and extended survival, effects potentiated by the BCL2 inhibitor venetoclax (**Figures 7H-7I**). Prostratin-treated tumors exhibited increased percentages of cleaved CASP3-positive cells and decreased percentages of Ki-67-positive cells, effects enhanced by venetoclax (**Figures 7J-7K**). Importantly, prostratin induced CASP8/CASP3/GSDME cleavage, particularly when combined with venetoclax (**Figure 7L**). Of note, prostratin remained effective against *GSDME* KO ST88-3 tumors (**Figures S15A-S15C**), increasing cleaved CASP3-positive cells while reducing Ki-67-positive cells (**Figures S15D-S15E**). The activated CASP8/CASP3 axis in *GSDME* KO tumors indicated prostratin could trigger apoptosis in the absence of GSDME (**Figure S15F**), demonstrating that GSDME is not required for the therapeutic efficacy of prostratin in NF1-deficient cancers. In contrast, prostratin showed no therapeutic effects in NF1-sufficient MPNST-724 xenografts (**Figures S16A-S16C**), with no significant changes in percentages of Ki-67-positive cells or pyroptotic/apoptotic cell death markers (cleaved CASP8/CASP3/GSDME) in tumor tissue (**Figures S16D-S16F**). Together, these results demonstrate therapeutic efficacy of PKC agonism for selectively targeting NF1-deficient tumors *in vivo*.

## Discussion

Achieving cell-specific pyroptosis for disease treatment represents a promising yet elusive therapeutic goal. Using NF1-deficient cells relying on KRAS as a model, this study demonstrates that PKCδ agonism triggers pyroptosis specifically in NF1-deficient cells, thereby suppressing NF1-deficient tumor growth (**Figure S17**). These findings establish that genetic vulnerabilities arising from cancer mutations can be pharmacologically targeted to induce cancer cell-specific pyroptosis, with immediate implications for diverse malignancies harboring *NF1* mutations and providing a conceptual framework for targeting other oncogene-addicted cancers.

Considerable progress has been made in understanding pyroptosis executioners (e.g., GSDMD and GSDME) and their immediate upstream proteases (e.g., caspases, granzymes, and elastase).^1^ However, molecules further upstream that regulate pyroptosis remain largely undefined. While regulators of caspase/GSDMD-dependent pyroptosis have been identified in infectious diseases,^1, 54^ upstream regulators of caspase/GSDME-dependent pyroptosis remain elusive, despite its relevance to many diseases including cancer.^1^ Here, we identify PKCδ as a critical upstream regulator of CASP8/CASP3/GSDME-dependent pyroptosis, filling this critical knowledge gap. Moreover, previous work showed that PKCδ directly phosphorylates NLRC4^54^ to activate CASP1/GSDMD-dependent pyroptosis.^55^ Together, these findings establish PKCδ as a hub regulator orchestrating both CASP1/GSDMD- and CASP3/GSDME-dependent pyroptotic programs through distinct substrate specificity (NLRC4 and KRAS, respectively). This regulatory framework, wherein a single kinase orchestrates different pyroptotic programs via specific substrates, may extend to other caspase-dependent regulatory mechanisms and alternative cell death programs, with implications beyond cancer.

Our findings expand the current RAS paradigm that defines RAS-GDP as “inactive” and RAS-GTP as “active,” because canonical RAS effectors including RAF proteins preferentially bind RAS-GTP.^18, 56^ In contrast, we found that KRAS-GDP preferentially interacted with CASP8 compared to KRAS-GTP, promoting CASP8 activation and cell death, thereby establishing CASP8 as a KRAS-GDP effector. This finding is consistent with evidence that the yeast Ras-like GTPase Bud1-GDP associates with Bem1 to regulate cell polarity,^57^ that Ran-GDP binds nuclear import factor p10 to modulate nucleocytoplasmic transport,^58^ and that phosphorylated RacE-GDP—where RacE is the *Dictyostelium* homolog of human RhoA—directly activates mTORC2 toward AKT to regulate cell migration,^59^ suggesting that phosphorylation-dependent activation of GDP-bound GTPase may represent a conserved regulatory mechanism across species and GTPase families. The functional relevance of the GDP-bound state is further underscored by the observation that multiple compounds and peptides preferentially bind GDP-bound GTPases, including KRAS-GDP.^60–63^ Intriguingly, KRAS-GTP and KRAS-GDP utilize distinct effectors to regulate opposing cellular fates: KRAS-GTP drives proliferation and transformation through canonical effectors, whereas KRAS-GDP promotes cell death via CASP8, revealing effector-dependent functional specialization based on nucleotide-bound states. Given the many cellular roles of KRAS,^18^ defining additional functional KRAS-GDP effectors and exploring whether this paradigm extends to other RAS paralogs will be of interest. This expanded view of RAS-GDP functionality provides new insights into RAS biology and suggests novel therapeutic avenues.

Our discovery that PKCδ phosphorylates KRAS to suppress its activity identifies a promising therapeutic strategy for RAS-driven cancers. Current FDA-approved KRAS(G12C) inhibitors are constrained by two limitations: they target only a specific mutation subtype and inevitably encountering acquired resistance.^64, 65^ PKC agonism offers a mechanistically distinct approach that inactivates KRAS through dual phosphorylation at S39 and S181, simultaneously accelerating GTP hydrolysis and impeding GTP reloading, thereby trapping KRAS in an inactive GDP-bound state regardless of mutation type. We demonstrate the efficacy of this approach in KRAS-mutant, NF1-deficient cancers, including triple-negative breast cancer (MDA-MB-231) and colorectal cancer (HCT-116) with G13D mutations (**Figures 6, S13**). Together with prior studies demonstrating prostratin efficacy in KRAS-driven pancreatic models,^24^ where KRAS mutations are universal, these findings collectively suggest PKC agonism may represent a pan-KRAS suppression strategy with the potential to address resistance mechanisms that plague current KRAS inhibitors, including compensatory NF1 loss.^64, 65^ Regarding the form of cell death induced by PKC agonism, an earlier study reported apoptosis in KRAS-mutant tumors based on CASP3 biosensor activation;^32^ however, this was subsequently revised to autophagic cell death in a follow-up study, as direct CASP8 cleavage was not detected following PKC agonist treatment.^31^ In contrast, our study establishes a mechanistically distinct outcome—CASP8/CASP3/GSDME-dependent pyroptosis—specifically in NF1-deficient cells, supported by direct biochemical evidence of caspase cleavage and gasdermin activation.

Our data establish that dually phosphorylated KRAS-GDP promotes CASP8 activation through competitive displacement of inhibitory BCL2 from a shared interface. BCL2 is well-known to regulate intrinsic apoptosis through BAX/BAK/tBid antagonism^35, 45^ and also physically sequesters CASP8, suppressing its activation; BCL2 inhibition with venetoclax releases this suppression.^46–48, 66^ Building on this mechanism, we demonstrate that KRAS competes with BCL2 to reduce BCL2-CASP8 interaction and promote CASP8 activation, an effect dependent on prostratin-induced dual phosphorylation of KRAS at S39 and S181. The BCL2-CASP8 interface residues we identified—K148/R149 in CASP8—are proximal to those previously described (V150/L157 in CASP8),^46^ supporting the structural basis of this interaction. Together, these findings establish phosphorylated KRAS as a necessary cofactor that initiates CASP8 activation by relieving BCL2-mediated suppression, although the full activation complex architecture remains to be characterized in future work. Regarding whether CASP8 activation in this context involves death receptor signaling, prior studies have shown that the PKCδ-dependent proapoptotic agent etoposide induces CASP8 activation in FADD-knockout cells without receptor-induced death-inducing signaling complex (DISC) formation,^67, 68^ demonstrating that PKCδ-driven CASP8 activation can proceed through death receptor-independent pathways. By analogy, given the shared PKCδ dependency, non-canonical receptor-independent pathways likely contribute similarly to prostratin-induced CASP8 activation in NF1-deficient cells.

Using prostratin, we define conditions for when and how to use PKC therapeutics in cancer,^69^ demonstrating anti-tumor activity of PKC agonism in NF1-deficient tumors but not NF1-sufficient tumors. Furthermore, selective PKCδ activation replicates the effect of prostratin, triggering the pyroptosis pathway (CASP8/CASP3/GSDME) in NF1-deficient cells. Intriguingly, loss-of-function mutations in PKCδ and CASP8, and gain-of-function mutations in KRAS, each independently cause autoimmune lymphoproliferative syndrome with defective cell death,^22, 69^ underscoring the functional relevance of our identified PKCδ/KRAS/CASP8 pathway. Identifying PKCδ as the specific PKC isozyme mediating these effects provides rationale for developing PKC isozyme-specific agonists for cancer treatment. Given the wide tissue expression of PKC isozymes and their differential, even opposing, roles in homeostasis and disease,^69^ isozyme-specific targeting could potentially reduce side effects and optimize therapeutic efficacy. For instance, PKCα/γ-activating agents may harm the central nervous system, as aberrant PKCα/γ activation due to germline mutations induces neurodegenerative diseases.^22^

The therapeutic implications of PKCδ agonism in NF1-deficient cancer extend beyond direct cytotoxicity. PKCδ agonism majorly induces pyroptosis, a form of immunogenic cell death that releases damage-associated molecular patterns and inflammatory cytokines to augment antitumor immunity.^3, 4^ Beyond tumor cell killing, PKC agonists directly enhance immune cell functions, including improved antigen presentation and reversal of T cell exhaustion.^70, 71^ This dual capacity distinguishes PKCδ agonism from MEK inhibition, which induces cytostatic senescence without immune activation.^11^ These immune-modulatory properties position PKC agonists as potential agents for converting immunologically “cold” tumors into “hot” tumors, with particular relevance for NF1-deficient tumors that resist immune checkpoint inhibitors due to impaired antigen presentation^72^ or poor immune cell infiltration.^73^ Validation in immunocompetent preclinical models and systematic evaluation of synergies with immune checkpoint inhibitors and other immunotherapies represent a critical next step of developing PKC agonism-based therapeutics for NF1-deficient cancers.

In summary, this work advances our understanding of NF1/RAS biology, provides insights into cell death regulation and therapeutic applications of PKC agonism, and demonstrates a pharmaceutical strategy for selective pyroptosis manipulation based on cell-specific molecular abnormalities.

## Limitations of the study

First, NF1 loss occurs in diverse tumors (including melanoma, glioma, and leukemia^74–76^), systematic studies beyond Schwann cells are needed to validate KRAS dependency, although prostratin induced pyroptosis in multiple NF1-deficient tumor types. Second, the mechanism underlying the RAS paralog-dependency switch from HRAS to KRAS upon NF1 loss warrants further investigation, likely reflecting differential regulation of RAS paralogs by NF1 and the distinct effector interactions and signaling outputs of HRAS versus KRAS when localized to different subcellular microdomains.^18^

## Materials and methods

### Reagents

A comprehensive list of reagents, including antibodies, commercial kits, bacterial strains, and cell lines used in this study, is provided in **Table S2.** Additional reagent specifications are detailed below.

### Experimental Model Details

#### Primary Mouse and Human Schwann Cell Derivation

##### Mouse Schwann Cell Derivation

*Nf1^+/-^* mice (Jackson Laboratory, #002646) were maintained through in-house breeding. Embryonic day 12.5 (E12.5) dorsal root ganglia (DRGs) were harvested for Schwann cell (SC) derivation, with E0.5 designated as the morning of the day when a vaginal plug was detected. Genotypes were confirmed for all derived SCs. Mouse SCs were cultured in DMEM (Thermo Fisher, #11965118) supplemented with 10% FBS, 1X penicillin-streptomycin, β-heregulin (10 ng/mL; Peprotech, #100-03), and forskolin (2 µM; Cayman Chemical, #11018) at 37°C with 7.5% CO_2_. Primary mouse Schwann cells from passages 2-3 were utilized for all experiments to ensure experimental consistency, as extended cultivation of these cells is not feasible in the specified culture medium.

##### Human Schwann Cell Derivation

Human SCs were isolated from plexiform neurofibroma (pNF) specimens surgically removed from NF1 patients. Tumor specimens were pathologically confirmed and obtained under a Waiver for HIPAA Authorization and Informed Consent (Mayo Clinic, IRB approved). Fresh tumor tissue was mechanically dissociated into ≤2 mm fragments and enzymatically digested in L-15 medium containing collagenase type I (0.5 mg/mL; Worthington Biochemical, #LS004196) and Dispase II (0.5 mg/mL; Sigma-Aldrich, #04942078001) for 1 hour at 37°C with orbital shaking (180 rpm). The cell suspension was filtered through a 40μm strainer to remove debris and treated with ACK lysis buffer (Thermo Fisher, #A1049201) to eliminate red blood cells. Cells were collected by centrifugation (500g, 4 minutes), washed once with ice-cold PBS, and plated. Human neurofibroma-derived SCs were seeded on Matrigel & Poly-L-lysine coated culture vessels and maintained in DMEM (Thermo Fisher, #11965118) supplemented with 10% FBS, 1X penicillin-streptomycin, IGF-1 (100 ng/mL; Peprotech, #100-11), β-heregulin (10 ng/mL; Peprotech, #100-03), and forskolin (2 µM; Cayman Chemical, #11018).

#### Schwann Cell Sphere Culture

Mouse SC spheres were generated as previously described.^1^ DRGs isolated from E12.5 mouse embryos were enzymatically dissociated with 0.25% Trypsin (Thermo Fisher, #25200056) for 3-5 minutes at 37°C, followed by mechanical dissociation using narrow-bore pipettes. The cell suspension was filtered through a 40μm strainer and centrifuged at 400g for 4 minutes. The resulting cell pellet was resuspended and cultured in ultra-low attachment 24-well plates (Fisher Scientific, #07-200-602) in serum-free DMEM/F12 medium supplemented with 1X N2 supplement (GIBCO, #17502048), EGF (20 ng/mL; R&D systems, #236-EG-200), and bFGF (20 ng/mL; Peprotech, #450-33). For human SC sphere formation, neurofibroma-derived cells were cultured in serum-free DMEM/F12 medium supplemented with 1X N2 supplement, IGF1 (100 ng/mL), EGF (20 ng/mL), and bFGF (20 ng/mL).

For sphere passaging, cultures were centrifuged at 400g for 3 minutes and dissociated with 0.05% Trypsin (Thermo Fisher, #25300054) for 5 minutes at 37°C. Single-cell suspensions were generated using narrow-bore glass pipettes and plated at 2,000-3,000 cells per well in 24-well plates. The culture medium was refreshed every 3-4 days.

For drug treatment studies, SC spheres from passages 2-3 were used, with three to four replicates per experiment. The treatment was started on day 0 after plating, and sphere numbers per well were counted using an inverted phase contrast microscope seven days post-treatment.

#### Established Cell Lines

All cell lines were routinely tested for mycoplasma contamination and authenticated through morphological assessment and growth characteristics according to ATCC standards (https://www.atcc.org) and the original derivation paper.^1^ The iHSC1λ cell line, malignant peripheral nerve sheath tumor (MPNST) cell lines (STS-26T, MPNST-724, ST88-14, ST88-3, S462, and NF90.8), and the Lenti-X 293T cell line were maintained at 37°C in a 5% CO_2_ atmosphere. These cells were cultured in a DF medium consisting of DMEM supplemented with 10% heat-inactivated fetal bovine serum (FBS), 1X penicillin-streptomycin, 2 mM L-Glutamine, and 1mM sodium pyruvate. Breast cancer and other cancer cell lines were maintained at 37°C in a 5% CO_2_ atmosphere, with the following media specifications: SK-MES-1: EMEM medium (ATCC, #30-2003) supplemented with 10% FBS; MCF-7, MDA-MB-468, MDA-MB-231, and HCT-116: DMEM medium (Thermo Fisher, #11965092) supplemented with 10% FBS. Cells were maintained with regular medium changes every 3-5 days and passaged using 0.05% trypsin-EDTA when reaching 70-80% confluency, with split ratios ranging from 1:4 to 1:10.

#### Genetically Modified Cell Lines

##### a. CRISPR-Based Modifications

CRISPR/Cas9 or CRISPR/Cpf1 was used for gene knockout: *NF1* in iHSC1λ cells, *CASP8* in ST88-3 and S462 cells, *GSDME* in iHSC1λ, S462, ST88-3, and MDA-MB-231 cells.

CasRx was employed for gene knockdown: *H-/N-/K-RAS* in iHSC1λ cells, *PKCδ/ε/η* and *KRAS* in S462 and ST88-3 cells.

##### b. Transgene expression

Transgenic expressions were implemented via either PiggyBac transposon system transfection or lentivirus infection. The following constructs were expressed:

In WT and/or *NF1* KO iHSC1λ cells: GFP, GFP-NF1-WT, or GFP-NF1-R1276P; Flag-HRAS/NRAS/KRAS (with S17N mutation); mRuby2-KRAS and PKCδ-GFP co-expression; tetO3G-GFP, tetO3G-PKCδ-CA (catalytic domain), or tetO3G-PKCδ-KD (kinase-dead, K378R); GFP-KRAS mutants (WT and phosphomutant S39A, S181A, or S39A/S181A) with CasRx-KRAS; GFP-KRAS-WT/-EE/-T35A cell lines. In ST88-3 *and* S462 cells: tetO3G-GFP, tetO3G-PKCδ-CA, or tetO3G-PKCδ-KD cell lines; GFP-KRAS mutants (WT and S39A/S181A) with CasRx-KRAS; mRuby2-KRAS and PKCδ-GFP co-expression; GFP-KRAS-WT/-T35A/-EE cell lines; GFP-KRAS-WT/-S39E/-S181E/-EE for immunostaining; GFP-KRAS-WT and CASP8-FL-D374A/D384A-mRuby2 co-expression; Flag-GFP or Flag-BCL2. In HEK293T cells: HA-StrepII-tag-KRAS-WT/-S39E/-S181E/-EE and HA-Nanoluc for protein expression; Flag-BCL2-ΔTM (removal of C-terminal signal peptide) for protein expression; GFP-KRAS-EE and CASP8 mutants (FL, Nter, Cter)-halotag co-expression; CASP8-FL-halotag and GFP-KRAS-WT/-T35A co-expression; CASP8-Cter-halotag (or Halotag) and GFP-KRAS-WT/-T35A co-expression; Flag-BCL2-ΔTM and CASP8 mutants (FL, Nter, Cter)-halotag co-expression. Additional modified cell lines: *GFP-KRAS-G13D* transgenic expression in MDA-MB-231 cells, and Flag-GFP or Flag-BCL2 expression in ST88-3 and S462 cells.

Modified cells were cultured in the same media as the parental cell line. Knock-out isogenic cell lines were cloned by limited dilution assays, knock-down cell lines enriched by drug selection (at least one week) or flow-sorting (post-transfection 72-96h), and transgene transduced cells were enriched through drug-selection (e.g., puromycin, zeocin, hygromycin) for at least one week and then maintained as the parental cell line.

#### Animal Studies

Animal studies were conducted in accordance with protocols (#20180043, #20210035, #20180103, #20210057) approved by the Institutional Animal Care and Use Committee (IACUC) at Cincinnati Children’s Hospital Medical Center (CCHMC). All mice, including genetically engineered mice on a mixed 129S/C57BL/6J (129/Bl/6) background and immunodeficient NSG mice, were housed and maintained under specific pathogen-free (SPF) conditions with a 14-hour light/10-hour dark cycle.

##### Genetically Engineered Mouse Model

As previously described,^3^ *Dhh-*Cre mice (Jackson Laboratory, #012929) and *Nf1*-flox (*Nf1*^fl/+^) mice (Jackson Laboratory, #017639) on a mixed 129/Bl/6 background were obtained and bred at CCHMC. Genotyping was performed by PCR, as detailed previously.^3^ For treatment studies, age-matched male and female *Dhh-Cre*, *Nf1*^fl/fl^ littermates were randomized and enrolled at seven months of age, coinciding with complete plexiform neurofibroma penetrance. Two-month treatments were administered by intraperitoneal injection and included vehicle (1% ethanol in 1x PBS) and prostratin (0.8 mg/kg in 1% ethanol/1xPBS).

##### Xenograft Studies

*NF1* KO iHSC1λ Cell Xenografts

Female NSG mice (Jackson Laboratory, #005557), aged 2-3 months, were obtained from CCHMC’s Comprehensive Mouse and Cancer Core. *NF1* KO iHSC1λ cells (2 × 10^6^) expressing *CasRx-NT*, *CasRx-HRAS*, or *CasRx-KRAS* were suspended in N2 medium: Matrigel (ratio of 1:2) containing β-heregulin (200 ng/mL) and injected subcutaneously into the right flank. Doxycycline chow was administered beginning three days post-injection and continued until study endpoint (tumor volume > 2,000 mm^3^, tumor ulceration, or nine months post-injection). Tumor volume was monitored biweekly for the first three months, then weekly thereafter. Mice were euthanized at the study endpoint for tissue analysis.

##### MPNST Xenografts

MPNST cells (3 × 10^6^; parental ST88-3, *GSDME* KO ST88-3, or MPNST-724) were suspended in N2 medium: Matrigel (1:3) supplemented with bFGF (100 ng/mL) and IGF1 (500 ng/mL), then injected subcutaneously into the right flank of female NSG mice (2-3 months old). Tumor measurements began on day 10 post-injection and continued every other day. Treatment was initiated when tumors reached approximately 200 mm^3^. Treatment groups included: vehicle (1% ethanol in 1x PBS, 60% Phosal-50 propylene glycol (PG) + 30% polyethylene glycol 400 (PEG400) + 10% ethanol), prostratin (1.25 mg/kg, intraperitoneal injection), venetoclax (25mg/kg, dissolved in 60% Phosal-50 PG + 30% PEG400 + 10% ethanol, oral gavage), and the combination (prostratin + venetoclax). Tumor volume (*V*) was calculated using the formula *V* = π/6 × *L* × *W*^2^, where L is length and W is width. Mice were euthanized when tumors reached 2,000 mm^3^ for tissue collection.

#### Vector Construction

All plasmids used for this study were confirmed by DNA sequencing at DNA sequencing and Genotyping Core at CCHMC.

##### CRISPR-Based Gene Knockout Vectors

Gene knockouts were performed using either the PX459 CRISPR/Cas9 vector (Addgene, #62988) or CRISPR/LbCpf1 vector (Addgene, #74042). sgRNA sequences were inserted following Addgene protocols (sequences listed in **Table S3**).

##### CasRx-Based Gene Knockdown Vectors

The CasRx lentiviral vector was constructed by inserting CasRx-2A-EGFP from pXR001:EF1a-CasRx-2A-EGFP vector (Addgene, #109049) into lentiCRISPR v2 (Addgene, #52961). For constitutive CasRx knockdown experiments, the EF1a promoter was replaced with the CAG promoter from pPB-CAG-MCS-IRES2-Puro vector (System Biosciences, gifted by Dr. Qi-Long Ying’s laboratory at the University of Southern California). For inducible CasRx knockdown, the EF1a promoter and puro resistance cassette were replaced with the tetO promoter and tetO3G-P2A-puro synthesized by Integrated DNA Technologies (IDT). Gene-specific knockdown spacer sequences were inserted into CasRx vectors using Esp3I (New England Biolabs, # R0734S), as described in **Table S3**.

##### Inducible Expression Vectors

The following tetracycline-inducible lentivectors were used for inducible gene knockdown and transgenic expression: *tetO-CasRx-T2A-EGFP*, *tetO*-*GFP-KRAS*, *tetO*-*PKCδ-CA*, *tetO*-*PKCδ-KD, tetO-CASP8-WT,* and *tetO*-*CASP8-C360S*.

##### Constitutive Expression Vectors

For constitutive lentiviral expression, *CAG-mGFP-P2A-Flag-MCS-IRES-Puro/Zeocin/Hygromycin* elements (System Bioscience) were inserted into lentiCRISPR v2 (Addgene, #52961).

##### Gene Cloning and Mutagenesis

Target genes were amplified by PCR using either gBlock gene fragments (synthesized by IDT) or iHSC1λ cDNA templates and subsequently cloned into expression vectors. Site-directed mutagenesis was performed using PCR. All mutagenesis primer sequences are listed in **Table S3**.

#### Gene Delivery

Gene delivery was achieved via transposon integration or lentiviral infection. Stable cell lines were established through drug selection or fluorescence-activated cell sorting (FACS). FACS was performed using the Bigfoot Spectral Cell Sorter (ThermoFisher) at CCHMC Research Flow Core.

##### Transposon-Mediated Transfection

For transposon delivery, PiggyBac mobile transposase (System Biosciences, gifted from Dr. Ying’s laboratory at the University of Southern California) and transposon vectors were mixed with polyethylenimine (1 mg/mL; Sigma Millipore, #919082) at a 1:3 ratio.

##### Lentiviral Production and Infection

Lentivirus was produced in Lenti-X cells (Takara, #632180) by co-transfection of psPAX2, VSVG, and transfer vectors containing gene expression cassettes according to the Addgene Lentivirus protocol. Viral supernatants were concentrated using Lenti-X™ Concentrator (Takara, #631231). Infection efficiency was enhanced by adding polybrene (8 μg/mL; Sigma Millipore, #TR-1003-G) to the virus-medium mixture during overnight incubation.

##### Electroporation of iHSC1λ Schwann Cells

For iHSC1λ cells resistant to conventional transfection methods, the Neon® Electroporation System (Thermo Fisher) was employed. Cells (2 × 10^5^) were washed with cold PBS and resuspended in 10 μL electroporation buffer R containing concentrated plasmid DNA (1.5 μg). These electroporation parameters (Pulse voltage: 1360 mV; Pulse width: 10 ms; Pulse number: 3) were used. The culture medium was refreshed the following morning, and drug selection was started 24 hours post-electroporation. Stably transfected cell lines were used for subsequent experiments.

#### RAS Pull-down Assays

Active RAS paralog levels were assessed using the RAS Activation Assay kit (Cell Biolabs, STA-400) in conjunction with RAS paralog-specific antibodies.^2^ Cells were grown to 80-90% confluency in 10 cm dishes and serum-starved overnight in DMEM/F12 medium containing 0.5X N2 supplement (GIBCO, #17502048). Cells were then stimulated for 5 min with 10% FBS plus β-heregulin (20 ng/mL; Peprotech, #100-03). Stimulated cells were washed with ice-cold PBS and lysed in 500μL 1x assay/lysis buffer (25 mM Tris-HCl pH 7.5, 150mM NaCl, 1% NP-40, 10mM MgCl_2_, 1mM EDTA, and 2% Glycerol) supplemented with 1x Halt™ protease/phosphatase inhibitors (Thermo Fisher, #78442). Lysates were kept on ice and gently triturated with large-orifice pipette tips every 10 minutes for 30 minutes, then cleared by centrifugation (17,000g, 10 minutes, 4°C). Five percent of the supernatant was kept as input. The remaining supernatant was incubated with RAF1-RBD agarose beads (30 μL) for 1-2 hours at 4°C with end-to-end rotation. Beads were washed four times with assay/lysis buffer containing protease/phosphatase inhibitors. Bound proteins were eluted in 60 μL of 2× SDS loading buffer containing 1 mM DTT, heated at 95°C for 5 minutes, and centrifuged (14,000 g, 30 seconds). Supernatants were analyzed by immunoblotting using RAS paralog-specific antibodies (**Table S2**).

#### Cell Viability and Proliferation Assay

##### CCK-8 Viability Assay

Cell viability was measured with the cell counting kit-8 (CCK-8) (APExBIO, #K1018) absorbance assay using the FlexStation® 3 Multi-Mode Microplate Reader (Molecular Devices). Cells (600-2,000 per well, depending on cell lines used) were seeded in 96-well flat-bottom plates (Corning, #353072) with three to five biological replicates (n=3-5). Test compounds were added to cells the following morning. 72-96 hours later, 10µL CCK-8 stock solution was added to cells in 96-well plates, and the absorbance was read at 450nm/650nm (650nm was used as the background signal) following the manufacturer’s instructions. Each condition included three to four biological replicates in 96-well plates, and the results represent the mean ± SEM of multiple replicates, expressed as a percentage of the control group after normalization. For growth curve studies, measurements were taken at 0, 24-, 48-, 72-, and 96 hours post-plating, and the plate was read after 1-2 hours incubation (same duration for different time points) in the 37°C incubator. For endpoint studies, the CCK-8 stock solution was added to cells 72-96 hours post-plating, and plates were read after 1-2 hours of incubation in the 37°C incubator.

##### EdU Proliferation Assay

For cell proliferation analysis, 2 × 10^4^ cells were seeded in triplicate in 24-well plates. After attachment, cells were incubated with 10 µM 5-ethynyl-2′-deoxyuridine (EdU) (Cayman Chemical, #20518) for 3 hours at 37°C. EdU incorporation was detected using the Click-iT™ EdU Imaging Kit with Alexa Fluor™ 594 (Thermo Fisher, #C10086) according to the manufacturer’s protocol.

#### Anchorage-Independent Soft Agar Assay

A base layer of 0.75% agar (Sigma, #A1296) in cell culture medium (DF medium with 10% fetal bovine serum) was prepared and allowed to solidify in 6-well plates. Cells (1.0-2.0×10^4^ per well in six-well plates) were suspended in 0.4% agarose (Thermo Fisher, #16500100) in cell culture medium and plated in triplicate over the base layer. For drug treatment experiments, compounds were added to the agarose cell suspension before plating. Cells were supplemented with 200 µL fresh culture medium with the compound every three days. For iHSC1λ cells, the culture medium was further supplemented with 10 ng/mL β-heregulin. After approximately four to five weeks, colonies were stained overnight with nitroblue tetrazolium chloride (1 mg/mL in 1× PBS). Images were acquired using an inverted microscope (Nikon) equipped with a digital camera.

#### Lactate Dehydrogenase (LDH) Release Assay

LDH levels in the cell supernatant were measured with the LDH-Glo™ Cytotoxicity Assay kit (Promega, J2380). Following vehicle/prostratin treatment, the supernatant was collected using 1x LDH storage buffer and properly diluted (100x to 400x) according to the manufacturer’s guideline, and the LDH release was quantified using the FlexStation® 3 Multi-Mode Microplate Reader with luminescence as the reading mode. The wells containing medium only, without any cells, were used as the background for correction. The percentage of LDH release was calculated based on the maximum LDH releases from cells subjected to the same treatment and lysed with 0.5% Triton X-100 at 37°C for 10 minutes.

#### Time-lapse Microscopy

Cell death kinetics were assessed with the Incucyte SX5 live-cell analysis system (Sartorius). Cells were plated at a density of 3,000-4,000 cells per well in 0.1% gelatin-coated 96-well chimney plates (Greiner, #655090) with four biological replicates and incubated overnight at 37°C with 7.5% CO_2_. Following overnight culture, cells were treated with vehicle or prostratin (2-4 μM) in the Opti-MEM medium (Thermo Fisher Scientific, #51985034) supplemented with 8% fetal bovine serum and 0.5 mM CaCl_2_ (to allow for Annexin-V binding). To quantify different types of cell death, the medium was supplemented with fluorescent dyes as indicated: SYTO-24 (Thermo Fisher, #S7559; 10nM, cell nuclear stain), SYTOX Orange (Thermo Fisher, #S11360; 10nM, pyroptotic/dead cell nuclear stain), and Annexin V-NIR (Sartoris, #4768; 1:3200 dilution, apoptosis/pyroptosis marker). Plates were maintained in the Incucyte SX5 system within a humidified incubator at 37°C with 7.5% CO_2_ throughout the 24-hour treatment period. Automated imaging was performed every 2 hours using green, orange, and near-infrared (NIR) fluorescence channels with a 10x objective lens. Four non-overlapping fields per well were captured at each time point. Image analysis and quantification were performed using Incucyte Analysis Software to determine cell death kinetics.

#### Pull-down Assays with Affinity Tag

Pull-down assays were performed using nanobody-based magnetic beads: GFP-trap (Proteintech, #gtma), RFP-trap (Proteintech, #rtma), halo-Trap (Proteintech, #otma), and anti-DDDDK tag magnetic beads (Abclonal, #AE126), following manufacturers’ protocols.

Cells expressing affinity-tagged proteins were cultured in 10 cm dishes, washed twice with ice-cold PBS, and lysed with 300 μL lysis buffer (20 mM Tris-HCl pH 7.5, 150 mM NaCl, 5 mM MgCl_2_, 0.5% NP-40) supplemented with Benzonase (Sigma, #E1014) and protease/phosphatase inhibitors (Thermo Fisher, #78442). Lysates were incubated on ice for 30 minutes with gentle trituration using wide-bore pipet tips every 10 minutes. After clearing by centrifugation (18,000 g, 20 min, 4°C), 5% of the supernatant was kept as input. The remaining lysate was transferred to a fresh tube and incubated with 25 μL pre-washed magnetic beads for 1-2 hours at 4°C with end-over-end rotation.

Beads were collected using a DynaMag-2 magnetic rack and washed 3-4 times with washing buffer (20 mM Tris-HCl pH 7.5, 150 mM NaCl, 5 mM MgCl_2_, 0.05% NP-40) containing protease/phosphatase inhibitors. Bound proteins were eluted directly in 50 μL of 2× SDS loading buffer supplemented with 1 mM DTT.

#### Phosphoproteomics

To identify prostratin-induced phosphorylation sites in KRAS4B, immunoprecipitation followed by phosphoproteomics analysis was performed. ST88-3 cells stably expressing GFP-KRAS4B-WT were serum-starved in DMEM containing 0.25% serum for 6h and then treated with either DMSO (vehicle control) or prostratin (4 μM) for 40 min. Cells were lysed in lysis buffer (25 mM Tris-HCl pH 7.5, 150 mM NaCl, 10 mM MgCl_2_, 1% NP-40) supplemented with Benzonase (Sigma, #E1014) and protease/phosphatase inhibitors (Thermo Fisher, #78442). GFP-KRAS4B was immunoprecipitated using GFP-trap magnetic agarose beads (Proteintech, #gtma) according to the manufacturer’s instructions. Immunoprecipitated material was eluted by boiling in 2× SDS sample loading buffer at 95°C for 5-10 minutes and resolved by SDS-PAGE on 4-20% gradient polyacrylamide gels (Bio-Rad, ##4561094). Gels were stained with AcquaStain protein gel stain solution (Bulldog Bio, #AS001000) on an orbital shaker for 30 minutes at room temperature, followed by three washes with distilled water to remove background staining. Protein bands corresponding to GFP-KRAS4B (∼50 kDa) were excised and submitted to the Taplin Biological Mass Spectrometry Facility (Harvard Medical School) for in-gel digestion (trypsin and Asp-N), phospho-peptide enrichment, and liquid chromatography-tandem mass spectrometry (LC-MS/MS) analysis.

KRAS-S181 phosphorylation was not detected despite the use of both trypsin and/or Asp-N digestion strategies. This likely reflects two compounding technical limitations: first, the repetitive lysine-rich sequences flanking S181 (KKKKKKS[181]KTK) produces short, poorly resolved tryptic peptides that are difficult to assign uniquely by database searching; second, the adjacent hydrophobic post-translational modifications at this region, including farnesylation of C185 and palmitoylation, are known to interfere with phosphopeptide enrichment efficiency and electrospray ionization, reducing detection sensitivity by mass spectrometry.

#### Protein Expression and Purification

##### Mammalian-Derived Proteins

KRAS, BCL2 lacking its C-terminal membrane-anchoring domain (BCL2-ΔTM, residues 1-211), and CASP8 C-terminal domain (CASP8-Cter) were expressed and purified from mammalian HEK293T cells. The coding sequences for these proteins, cloned from cDNA of human iHSC1λ cells or IDT gBlock fragment, were cloned into lentiviral vectors and verified by DNA sequencing. Stable 293T cell lines were established through drug selection following transfection. Cells were expanded in 150mm dishes (50-80 μg protein yield per dish) to generate sufficient protein for *in vitro* studies.

Proteins were purified using Strep-tag®II affinity chromatography according to the manufacturer’s protocol. To purify proteins, cells were lysed in ice-cold buffer (20mM Tris-HCl pH7.5, 150mM NaCl, 5mM MgCl_2_, 0.5mM CaCl_2_) supplemented with biotin blocking solution (IBA Lifesciences, #2-0205-050), Benzonase (Sigma, #E1014), and protease/phosphatase inhibitors (Thermo Fisher, #78442). Lysates were cleared by centrifugation (18,000 g, 20 min) and filtered through 0.45μm PES Filters. Cleared lysates were applied to the Strep-Tactin®XT 4Flow® column (IBA Lifesciences, #2-5011-005), washed five times, and eluted with 1x BXT buffer (IBA Lifesciences, #2-1042-025). Proteins were buffer exchanged to remove biotin in the BXT buffer, concentrated using 10 kDa MWCO centrifugal filters, and quantified by Bradford assay (Thermo Fisher, #23246).

##### Bacterial Expression of RAF1-RBD proteins

GST-tagged RAF1-RBD (residues 1-149) was expressed in *E. coli* BL21(DE3) (New England Biolabs, #C2527H) using the pGEX-KG vector. Cultures were grown in LB medium with ampicillin (100 μg/mL) and induced at OD_600_ = 0.8 with 0.2 mM isopropyl β-D-1-thiogalactopyranoside (IPTG) overnight at 20°C. Cells were harvested and lysed by sonication in buffer B (25 mM HEPES pH 7.4, 1 M NaCl) containing 1 mM PMSF and 10 mM DTT. The protein was purified using GST-affinity chromatography (GSTrap FF, Cytiva) and eluted in 100 mM Tris-HCl pH 8.0, 150 mM NaCl, and 10 mM reduced glutathione. Final purification was achieved by size-exclusion chromatography (HiLoad 16/600 Superdex 200 pg, Cytiva) in 20 mM Tris pH 7.5, 100 mM NaCl, and 1 mM DTT. Proteins were concentrated using 30 kDa MWCO filters and quantified by Bradford assay.

##### Cell-free Protein Expression (CFPE) of CASP8 Proteins

CASP8 Proteins (full-length F122G/L122G/D374A/D384A and N-terminal F122G/L122G; hereafter, CASP8-FL and CASP8-Nter) were produced using the ALiCE® CFPE system (LenioBio, #AL00000001). Codon-optimized sequences with C-terminal 6His-Strep-tag®II were cloned into the Alice01 vector, and the sequence was confirmed. Expression reactions (50 μL) containing vector DNA (5 nM) and RNase inhibitor (Promega, #N2511) were incubated at 25°C for 48 hours with constant shaking (700 rpm). After centrifugation (10,000 g, 5 min, 4°C), proteins were purified from the supernatant using Strep-tag®II affinity chromatography.

##### Quality Control and Tag Removal

Protein purity (>90%) was assessed using Instant-band dye (EZBiolab, #PFS001P) with BSA standards (Biorad, # 5000206) and confirmed by immunoblotting. KRAS and CASP8 functionality was verified through pull-down and binding affinity assays. For tag removal, proteins containing HRV-3C protease sites were sequentially incubated with Halo-trap beads and HRV-3C protease overnight at 4°C, followed by GST-affinity magnetic bead treatment to remove the GST-tagged protease. The final products were buffer-exchanged, concentrated, and quality-controlled.

#### *In Vitro* Kinase Assay

The assay was performed as previously described.^5^ Reaction mixtures (25 μL) contained recombinant PKCδ (100 ng; Abcam, #ab60844), HA-KRAS (2 μg), and 1× kinase buffer (20 mM HEPES pH 7.4, 150 mM NaCl, 10 mM MgCl2, 0.1 mM ATP) supplemented with protease inhibitors (ApexBio, #K1007) and PKC lipid activator (EMD Millipore, #20-133). For dephosphorylation controls, samples were treated with Lambda Protein Phosphatase (New England Biolabs, #P0753S) according to the manufacturer’s protocol. Reactions proceeded for 60-90 minutes at 30°C and were terminated by the addition of 5x SDS sample buffer followed by heat denaturation (95°C, 5 min). Phosphorylated proteins were analyzed by SDS-PAGE and detected using anti-phosphoserine and anti-phosphothreonine antibodies.

#### Nucleotide Exchange of KRAS

##### Nucleotide Exchange *In vitro*

Purified KRAS proteins were GDP-bound (due to GTP hydrolysis), as verified by RAF1-RBD pull-down and affinity binding assays. For GTP-loading as previously described,^3^ KRAS-GDP was incubated with 10-fold molar excess of GTPγS (Roche, #10220647001) and calf intestine phosphatase (CIP) agarose beads (Sigma, #P0762) in 1mL nucleotide exchange buffer (40 mM Tris pH 7.5, 200 mM (NH4)_2_SO_4_, 10 μM ZnCl_2_, 5 mM DTT) overnight at 4°C with rotation. Following exchange, the protein in the supernatant was purified by size-exclusion chromatography to remove excessive GTPγS. Successful GTPγS loading in KRAS was confirmed by RAF1-RBD pull-down and affinity binding assays.

##### Nucleotide Exchange in Cell Lysates

For nucleotide exchange (GDP/GTPγS loading) of GFP-KRAS in cell lysates, cleared lysates (500 μl) were incubated in GTPase loading buffer (20 mM Tris-HCl pH 7.4, 150 mM NaCl, 10 mM EDTA) containing either GDP (1 mM; Sigma, #7127) or GTPγS (1 mM) to convert KRAS into the GDP- or GTPγS-bound state. After incubation at 30°C for 40 min, the reaction was stopped by adding 2M MgCl_2_ to a final concentration of 15 mM. The equilibrated lysate was subjected to GFP-trap magnetic bead pull-down assays. The nucleotide-binding state of GFP-KRAS was confirmed by immunoblotting for RAF1 in the GFP pull-down fraction.

#### Binding Affinity Analysis

Direct protein-protein interaction affinities were measured using immobilization-free Microscale Thermophoresis and Spectral Shift assays.

CASP8 and GST-RAF1-RBD proteins were fluorescently labeled using the Protein Labeling Kit RED-NHS 2nd Generation (NanoTemper Technologies, # MO-L011). In brief, proteins (100 μM in 20μl) were incubated with a three-fold molar excess of dye for 30 minutes at room temperature in darkness. Excess dye was removed using the B-Column of the labeling kit. Final protein concentration (∼2 μM) and labeling efficiency (0.8-0.99) were determined by Nanodrop.

Thermophoresis measurements were performed using a NanoTemper Monolith NT.115 instrument. The system employs an infrared laser (IR-laser) coupled to the fluorescence light path via a dichroic mirror, focusing through shared optics into the sample fluid. The IR laser (≤24 mW) creates local heating within a 25 μm diameter region without compromising biomolecule integrity.^4^ Labeled GST-RAF1-RBD (0.025 μM), was incubated with 16 serial dilutions of GDP/GTPγS-loaded KRAS proteins in assay buffer (50 mM Tris pH 7.4, 150 mM NaCl, 0.05% Tween -20, 10 mM MgCl_2_, 10 μM GDP/GTPγS). Thermophoresis was monitored for 30 seconds at room temperature. Data from three or more independent experiments were normalized to ΔFnorm [‰] (10*(Fnorm(bound) - Fnorm(unbound))) and converted to fraction bound (0-1) by dividing by curve amplitude. Dissociation constants (Kd) were calculated using nonlinear regression (GraphPad Prism 9.2.1).

Spectral shift measurements were performed using a NanoTemper Monolith X instrument, which detects binding-induced spectral shifts at 650 nm and 670 nm under isothermal conditions.^5^ KRAS proteins were diluted to 20-40 μM in assay buffer (50 mM Tris-HCl pH 7.4, 150 mM NaCl, 0.1% Tween-20, 10 mM MgCl_2_, 10 μM GDP/GTPγS), while BCL2 proteins (BPS Bioscience, #50272) were diluted in the same buffer without MgCl_2_. Sixteen-point dilution series were prepared with the highest ligand concentrations of 5-20 μM and labeled CASP8 at 0.05-0.075 μM. Samples were equilibrated for 30 minutes before loading into capillaries and the Monolith X device. Measurements were performed at 25°C with three or more replicates. Data analysis was performed using MO.Control software (NanoTemper) with curve fitting and Kd calculations in GraphPad Prism 9.2.1.

#### GTPase Activity Assay

KRAS GTPase activity (intrinsic and GAP-stimulated) was measured using the GTPase-Glo™ Assay (Promega, #V7681) in 384-well plates as previously described.^3^ For intrinsic activity, GDP-bound KRAS proteins (WT, S39E, S181E, EE; up to 2 μg) at 16 concentrations were incubated with GTP substrate (1 μM) in 10 μL GTPase/GAP reaction buffer (50 mM Tris-HCl pH 7.5, 50 mM NaCl, 20 mM EDTA, 5 mM MgCl_2_) for 90 minutes at room temperature. GAP-stimulated assays included 200 ng GST-p120RasGAP (RASA1 GRD domain; BPS Bioscience, #100518-1).

Following the GTPase reaction, GTPase-Glo reagent (10 μL) was added and incubated with constant shaking for 40 minutes at room temperature to convert the remaining GTP to ATP. A detection reagent (20 μL) was then added and incubated for 5-10 minutes before measuring luminescence using a FlexStation® 3 reader (Molecular Devices). Each experiment was performed with at least four replicates, and at least three experiments were done for each KRAS protein.

For dose-response analysis, luminescence values were normalized to the maximum signal for each protein and fitted using nonlinear regression (4-parameter sigmoid, concentration as independent variable) with constraints (IC50 > 0, Top = 1, Bottom = 0). EC50 values (KRAS concentration producing 50% maximal GTP hydrolysis) were compared between proteins using the Extra sum-of-squares F Test (GraphPad Prism 9.2.1).

#### Nucleotide Dissociation Assay

As described previously,^3^ GDP-bound KRAS proteins (WT, S39E, S181E, EE; 500 nM) were incubated with fluorescent-labeled BODIPY FL-GDP (250 nM; Invitrogen) in buffer (20 mM Tris-HCl pH 7.5, 100 mM NaCl, 1 mM EDTA) at 25°C for 2 hours. MgCl_2_ was added to 5 mM final concentration, and baseline fluorescence was monitored for 10 minutes to ensure stability. Nucleotide exchange was initiated by adding GTPγS (5 μM; 10-fold excess) in the presence of 5 mM MgCl_2_.

Fluorescence intensity was monitored at one-minute intervals for 30 minutes using an EnVision Multimode Plate Reader (Perkin Elmer; excitation/emission: 485/535 nm). Data were baseline-corrected and normalized to initial fluorescence values. Dissociation rate constants (Koff) were determined using nonlinear regression with a one-phase exponential decay model (GraphPad Prism 9.2.1) from four or more independent experiments.

#### Protein Competition Assay *in vitro*

CASP8-FL Halotag fusion protein was prepared using the ALiCE® CFPE kit (LenioBio, #AL00000001), while HA-KRAS-EE, HA-KRAS-AA, and Flag-BCL2-deltaTM were purified from HEK293T cells. All proteins were purified using Strep-tag®II affinity chromatography and validated by instant-band gel imaging and immunoblotting.

For competition assays, CASP8-FL Halotag protein (50nM) and Flag-BCL2-deltaTM (500nM) were combined in 300μL assay buffer (50 mM Tris pH 7.4, 150 mM NaCl, 0.05% Tween-20, 10 mM MgCl_2_, 10 μM GDP/GTPγS) and incubated at room temperature for 90 minutes to form CASP8-BCL2 complexes. Next, varying concentrations of GDP-bound HA-KRAS-EE (0, 45, 180, 720nM) or GTPγS-bound HA-KRAS-AA (45, 180, 720nM) were added to the pre-formed CASP8-BCL2 complexes to assess competition between KRAS and BCL2 for CASP8 binding. After 30 minutes at room temperature, 5% of the reaction mixture was kept as input, and the remainder was incubated with Halo-trap beads (8 μL) to immune-precipitate CASP8-binding complexes. To minimize non-specific binding, Halo-trap beads were pre-blocked with 0.2% BSA overnight in low-protein binding Eppendorf tubes (Thermo Fisher, #901410). Following 60 minutes of incubation, beads were washed four times with the assay buffer containing 0.05% Tween-20. Bound proteins were eluted by boiling in 2× loading buffer (G-biosciences, #7867-701). Three independent experiments were performed.

#### Structure Prediction and Analysis

The KRAS-CASP8 complex structure was predicted using AlphaFold2^6^ implemented in ColabFold notebooks (ColabFold v1.5.5) on Google Colaboratory with default settings and Amber relaxation (parameters:msa_mode=mmseqs2_uniref_env, pair_mode=unpaired_paired, template_mode=pdb100, num_relax=5, model_type=alphafold2_multimer_v3, num_recycles=6, recycle_early_stop_tolerance=auto, relax_max_iterations=200, pairing_strategy=greedy, max_msa=auto, num_seeds=1). ColabFold differs from native AlphaFold2 primarily in its use of mmseqs2, which produces comparable results as reported by the ColabFold developers.^7^ For the prediction, protein sequences were concatenated with semicolon separators.

The BCL2-CASP8 complex was predicted with the Protenix platform (https://github.com/bytedance/Protenix), which implements the AlphaFold3 algorithm.^8,9^ The parameters for prediction were set as follows: “use_msa”: true, “atom_confidence”: true, “model_seeds”: [21448], “model_version”: “v1”, “N_sample”: 5, “N_step”: 200, “N_cycle”: 10.

All predictions were run with all five trained AlphaFold models. The resulting structures were evaluated for consistency, and the top-ranked model was selected for structure visualization and interface analysis. Prediction accuracy was assessed using AlphaFold’s confidence metrics: pLDDT (predicted Local Distance Difference Test), pTM (predicted Template Modeling), and ipTM (interface pTM) scores.^6,8^ Following ColabFold and AlphaFold recommendations, pTM and ipTM scores were used to rank predicted protein-protein complexes.^6–8^

For structure visualization, per-residue pLDDT scores (ranging from 0 to 100, with higher scores indicating better prediction) were converted to 100-pLDDT and stored in the B-factor column of PDB files. When displayed as pseudo-B-factors in Chimera-X,^10^ this transformation generates a light spectrum from red (lowest pLDDT) to blue (highest pLDDT).

#### Immunoblotting Analysis

Cultured cells were lysed directly in RIPA buffer (Teknova, #R3792) containing protease inhibitors, cleared by centrifugation (20,000 g, 15 minutes), and quantified by Bradford assay (Thermo Fisher, Thermo Fisher, #23246). For xenograft tumors, tissue was minced, weighed, and homogenized using a TissueRuptor (Qiagen, #9002755) in the RIPA buffer (500 μL per 100 mg tissue) supplemented with 10 mM EDTA and protease inhibitors. Tumor homogenates were sonicated and centrifuged, followed by concentration determination by Bradford assay.

Proteins (20-50μg) were denatured in 1× loading buffer (G-biosciences, #7867-701) at 95°C for 5-10 minutes and separated on 4-20% TGX precast gels (Biorad, #4561093, #4561095). Proteins were transferred to Immobilon®-P PVDF Membrane (Millipore, #IPVH00010) and blocked with 4% Blotting-Grade Blocker (Biorad, #1706404XTU) in 1xTBS for one hour at room temperature. For phosphorylation detection, PVDF membranes were fixed with 4% paraformaldehyde/0.01% glutaraldehyde^11^ for 35 minutes at room temperature, washed three times, and blocked with 4% BSA (Sigma, #A7030) for one hour.

Membranes were incubated overnight at 4 °C with primary antibodies (dilutions in **Table S2**). After washing three times (5 min each) with 1x TBS-T (0.05% Tween-20), membranes were incubated with horseradish peroxidase (HRP)-conjugated secondary antibodies (Cell Signaling Technology, #7077S, #7076S, 1:3000-1:5000 dilutions) for 1-2 hours at room temperature. Proteins were visualized on an Azure 600 Imaging system using the Immobilon Western Chemiluminescent HRP Substrate (EMD Millipore, #WBKLS0500).

#### Reverse Transcription and PCR

Total RNA was extracted with the Quick-RNA Miniprep Kit (Zymo Research, #R1054) and reverse transcribed with a high-capacity cDNA reverse transcription kit (Thermo Fisher, #4368813).

Conventional RT-PCR reactions (20μl) were performed with EconoTaq® PLUS GREEN 2X PCR Master Mix (VWR, #95024-006). Products (8μl) were analyzed by agarose gel electrophoresis after 30 cycles of amplification (except β-actin for 25 cycles) and imaged using an Azure 600 Imaging system.

qPCR was performed using Power SYBR™ Green PCR Master Mix (Thermo Fisher, #4367659) on a QuantStudio™ 6 Flex Real-Time PCR instrument with compatible 96-well or 384-well plates (Thermo Fisher). Default parameters for differential gene expression analysis were used, including melting curve analysis, to confirm product specificity. Relative expression levels were calculated using the ΔΔCt method with β-actin normalization. All PCR primer sequences are listed in **Table S3.**

#### TCGA Data Analysis

Epistasis analysis between NF1 and individual PKC genes (in total nine) was performed using the TCGA pan-cancer dataset (10,443 samples with mutation data) through cBioportal For Cancer Genomics (https://www.cbioportal.org/).^12^

#### Histological Analysis

Mice were perfused with ice-cold PBS, followed by overnight fixation in 4% paraformaldehyde at 4°C. After washing three times in PBS, mice were immersed in 30% sucrose solution overnight. Paraspinal cord tumors were then dissected, harvested, and embedded in Tissue-Tek® O.C.T. compound. For MPNST xenograft, immunodeficient NSG mice reaching the endpoint were euthanized, followed by perfusion, and tumors were removed, fixed in 4% paraformaldehyde overnight at 4°C, washed with PBS, and cryoprotected in 30% sucrose solution. Processed tumors were embedded in the O.C.T. compound. Serial sections (10-12 μm) were prepared for all tissue types, with a minimum of three slides containing nine sections per mouse.

Sections were fixed in 4% paraformaldehyde, permeabilized with 0.3% Triton X-100, and incubated with primary antibodies overnight at 4°C. Fluorescent secondary antibodies (Alexa®-488, -568, or -594; Thermo Fisher) were applied for 1-2 hours at room temperature. Nuclei were counterstained using VECTASHIELD® antifade mounting medium with DAPI (Vector Laboratories, #H-1900). Sealed sections were imaged using a Nikon C2+ Confocal microscope. Quantification was performed using CellProfiler 4.2.5 software^13^ (https://cellprofiler.org/) with the Cell/Particle counting pipeline to score the percentage of stained objects (https://cellprofiler.org/previous-examples).

#### Super-Resolution Microscopy

For super-resolution imaging, µ-Slide eight-well glass chamber slides (ibidi, #80807) were sequentially coated with Cell-Tak (Corning, #354240; 3.5μg/cm^2^) diluted in 0.1 M NaHCO_3_, followed by Laminin (Corning, #354232; 5μg/cm^2^), according to the manufacturer’s instructions. GFP-KRAS expressing cells were seeded at a density of 10,000-15,000 cells per well and treated with vehicle (DMSO; 0.1%) or prostratin (1μM) overnight (16-18h).

For CASP8 imaging, cells were transduced with a CASP8-C360S-HaloTag lentivirus (the C360S mutation renders CASP8 catalytically inactive, enabling stable expression without inducing cell death) for 60 h prior to seeding. Immediately before fixation, cells were incubated with Janelia Fluor 646 HaloTag Ligand (Promega, #HT1060; 200 nM) for 1 h at 37°C to label CASP8-C360S-HaloTag, followed by two washes with phosphate-buffered saline to remove unbound ligand.

Cells were fixed in 4% paraformaldehyde, permeabilized with 0.002% digitonin to preserve protein and organelle ultrastructure and incubated with primary antibodies overnight at 4°C. Fluorescent secondary antibodies (Alexa fluor®-568; Thermo Fisher) were applied for 1-2 hours at room temperature. Nuclei were counterstained using DAPI (Thermo Fisher, #D3571), and slides were mounted with non-hardening mounting medium (ibidi GmbH, #50001) prior to imaging.

Images were acquired on a Nikon AXR with NSPARC super-resolution microscope equipped with a four-line laser illumination system (405-nm, 488-nm, 561-nm and 640-nm) at the CCHMC Bio-Imaging and Analysis Facility (RRID:SCR_022628). Super-resolution images were acquired using the 8K galvo scanner in super-resolution mode with a 100X high NA oil immersion objective. Images were deconvoluted using the NIS Elements AI Suite (v6.5.20). Colocalization analyses were performed using the BIOP JACoP in Fiji (National Institute of Health, USA) with the following parameters: Threshold-Otsu (or Manual Threshold), Costes Block Size = 3 (for super-resolution images), and Costes Number of Shuffling = 200, and thresholded Manders Coefficients (thresholded M1) was computed. All the data passed Costes’ Significance Test (Costes P-Value > 0.95). For each protein pair analyzed, a minimum of five cells per replicate across at least two independent replicates (>=10 cells total) were quantified.

### Statistical Analysis

Data analyses were performed in Graphpad Prism (version 9.2.1). Sample sizes were not predetermined by statistical methods, and experiments were conducted without blinding. All *in vitro* experiments included at least three independent biological replicates (n>=3), and findings were confirmed across a minimum of two independent experiments or two different cell lines. The number of biological replicates (mice for *in vivo* studies) is specified in figure legends or figures. The equality of variances between groups was assessed before selecting statistical tests. For comparisons of two groups, data were analyzed using unpaired Student’s t-tests, with Welch’s correction applied when variances were unequal. For multiple comparisons (three or more groups), ordinary one-way ANOVA was used; for tumor growth kinetic curves, two-way ANOVA was used. Multiple comparisons among groups with unequal variances were analyzed using Brown–Forsythe and Welch ANOVA tests with Dunnett’s T3 correction. For two categorical variables (tumor-bearing versus non-tumor; mutant versus wild-type), Fisher’s exact test was used. For analyzing overall survival of tumor-bearing mice, the log-rank test was used to determine statistical significance. For binding affinity and RAS activity assays, statistical methods are described in previous sections. All data shown are reported as mean ± SEM, and error bars in figures represent SEM of biological replicates. *P*<0.05 was considered statistically significant, and the significance level was indicated with different numbers of asterisks. If 0.05<*P*<0.1, the *P*-value is shown; if P≥0.1, ns (non-significant) is indicated.

## Resource availability

### Lead contact

Further information and requests for resources should be directed to, and will be fulfilled by, the lead contact, Nancy Ratner.

### Materials availability

Cell lines and vectors generated in this study are available on request, but we may require a materials transfer agreement (MTA) and/or payment if there is potential for commercial application.

### Data and code availability

This paper does not report the original code. Any additional information required to reanalyze the data reported in this paper is available from the lead contact upon reasonable request.

## Acknowledgements

We thank Nicolas Nassar and Elliott Robinson (Cincinnati Children’s Hospital Medical Center, CCHMC), and all members of the Ratner laboratory for helpful discussion and feedback. Biplab Dasgupta (Emory University) and Nicolas Nassar kindly provided the breast cancer cell lines MCF-7 and MDA-MB-468, respectively. We thank Robert Hennigan (CCHMC) for technical assistance with protein purification and live-cell imaging with the Incucyte SX5, and Ronald Waclaw (CCHMC) for confocal microscopy training and technical support. We thank Matt Kofron and the Bio-Imaging and Analysis Facility (CCHMC; RRID:SCR_022628) for assistance with super-resolution imaging, Kenneth Quayle and the Research Flow Cytometry Core (CCHMC; RRID:SCR_022635) for flow sorting, and Victoria Summey and the Comprehensive Rodent and Radiation Facility (CCHMC; RRID:SCR_022624) for providing NSG mice for this study. We thank the Taplin Mass Spectrometry Facility at Harvard Medical School for performing the phosphoproteomics analyses, and NanoTemper Technologies for assistance with spectral-shift affinity assays. We thank Daniel Starczynowski and Nicolas Nassar (CCHMC) for critical reading of the manuscript. This work was supported by NIH R37 NS083580, NIH R01 NS120892, and a CancerFree KIDS award to N.R., NIH R01 CA278756 to Y.Z., and a Strauss Fellowship (2021), a CancerFree KIDS award (2024), and a Pardee transitional grant (2026) to L.H. The schematics were generated using https://www.Biorender.com.

## Author contributions

L.H. and N.R. conceived the project. L.H. drafted the manuscript and performed most of the experiments. L.H., Y.T.T., Y.L., Y.Z., and N.R. revised the manuscript. L.H., Y.Z., N.R., and Y.T.T. designed experiments. Y.L. conducted protein affinity and RAS nucleotide exchange assays and assisted with protein structure prediction and analysis. Y.T.T. contributed to protein purification. Y.T.T. and J.W. assisted in performing xenografts, and A.S. in drug administration. J.P. and T.R. helped with mouse tumor dissections and cell sorting. K.C.S. and J.W. generated primary mouse Schwann cells and human neurofibroma spheres, respectively.

